# Macaque Area LIP Reflects Confidence-Dependent Changes in Decision Policy

**DOI:** 10.64898/2026.04.27.721150

**Authors:** Mengyu Serene Tu, Miguel Vivar-Lazo, Christopher R. Fetsch

## Abstract

The process of forming a decision gives rise to an estimate of its quality or likelihood of success. This graded sense of confidence is important for guiding subsequent decisions, but a neural mechanism linking subjective confidence to formation of the next decision has not been identified. We trained rhesus monkeys to report a perceptual choice and simultaneous confidence judgment in a reaction-time (RT) motion discrimination task. Monkeys were more likely to repeat a rewarded choice when they reported low confidence on the previous trial, and showed greater changes in RT after a surprising outcome (low-confidence correct or high-confidence error). Neural activity in the lateral intraparietal area (LIP) encoded the previous trial’s choice and confidence more strongly after a low-confidence correct trial, and the strength of history encoding was correlated with a trial-by-trial estimate of choice bias. Ramping dynamics of the decoded decision variable also depended systematically on the conjunction of confidence and reward on the previous trial, in a manner that reflected individual differences between animals. The findings suggest that LIP not only reflects the formation of the current decision but could participate in confidence-guided learning via adjustments of the subsequent decision process.

## INTRODUCTION

We live in a changing and uncertain world where sensory information is often noisy and incomplete, and survival depends on learning from experience. The impetus to learn is so strong that humans and animals continually adjust their behavioral response to sensory input even when doing so is counterproductive, such as in psychophysical tasks where each trial is independent (from the experimenter’s perspective). The optimal strategy in such tasks is to make the decision based entirely on the sensory evidence in the present trial, yet subjects continue to be biased by the outcome of previous trials (Gold et al., 2008; Hwang et al., 2017; Mochol et al., 2021; Urai et al., 2017) even after extensive training (Gold et al., 2008; Mochol et al., 2021). In addition to history effects on choice bias, subjects often adjust the speed of their decisions following feedback (Beatty et al., 2021; Laming, 1979; Purcell & Kiani, 2016), often interpreted as a strategy to prevent future errors (Botvinick et al., 2001; Laming, 1979). In general, trial-history effects can be framed as adjustments of internal criteria and other parameters of the decision process, here collectively referred to as a decision policy.

Decision confidence, the subjective belief that a decision is correct, is crucial for optimizing learning from feedback (Drugowitsch et al., 2019; Lak et al., 2017; Mendonça et al., 2020). The logic is simple: a surprising outcome (high-confidence error or low-confidence correct choice) should drive faster learning because it suggests a change is warranted in the animal’s stimulus-response mapping or internal model. This idea can be formalized in the framework of reinforcement-learning (RL), where confidence may be considered as an ingredient in the computation of reward prediction error (RPE; Lak et al., 2020; Frömer et al., 2021). In one meta-analysis, mice, rats, and humans were more biased toward a previously rewarded choice when the previous trial was more difficult (Lak et al., 2020), consistent with confidence playing a role in RPE and hence learning rate. In another study, rats’ choice biases could be explained by learning from reward statistics in proportion to a model-based estimate of confidence (Mendonça et al., 2020). However, neither study attempted to measure subjective confidence directly, but instead relied on proxies related to stimulus strength and/or RT (Braun et al., 2018). Lastly, it has been suggested that confidence plays a role in adjusting the speed-accuracy trade-off for subsequent decisions (Desender et al., 2019; van den Berg et al., 2016). We therefore exploited a task design that measures choice, confidence, and RT on every trial (Vivar-Lazo & Fetsch, 2026), and trained macaque monkeys to perform the task to study the underlying neural mechanisms.

Although many frontoparietal and subcortical areas are likely involved, here we focus on the lateral intraparietal area (LIP) of the macaque posterior parietal cortex (PPC). When decisions are indicated with a saccadic eye movement, LIP populations represent an evolving decision variable (DV) that incorporates accumulated sensory evidence (Shadlen & Newsome, 2001; Steinemann et al., 2024) and predicts decision confidence (Kiani & Shadlen, 2009; Vivar-Lazo & Fetsch, 2026). Some neurons in LIP also carry information about trial history (Purcell & Kiani, 2016; Seo et al., 2009), as does the PPC in mice (Hwang et al., 2017; Morcos & Harvey, 2016) and rats (Akrami et al., 2018). However, it remains unknown whether LIP/PPC encodes confidence in a previous choice, and hence whether this region could mediate confidence-dependent adjustments of the decision process.

We hypothesized that LIP maintains or receives information about confidence on the previous trial, and that these ‘confidence history’ signals predict the adjustments of choice bias and RT in the current decision. Behaviorally, we found that the magnitude of trial-history effects was correlated with the monkey’s self-reported confidence on the previous trial, even when controlling for the objective stimulus strength. Consistent with the hypothesis, LIP population activity before and after stimulus onset encoded the previous trial’s outcome (choice and wager), and did so more strongly after a low-confidence rewarded trial. Moreover, single-trial instances of a decoded DV reflected confidence-dependent dynamics consistent with each animal’s distinct pattern of post-error RT. The results illuminate a key component of the distributed neural system mediating adaptive control of decision policy: the incorporation of a subjective belief in the correctness of a previous decision.

## METHODS

### Behavioral task and data acquisition

For the current study we reanalyzed a dataset described in previous work (Vivar-Lazo & Fetsch, 2026). Two rhesus monkeys (*Macaca mulatta*) were trained to perform a reaction-time motion discrimination task with peri-decision wagering (peri-dw). In each trial, the monkey viewed a dynamic random-dot stimulus and reported the net direction of motion with a saccadic eye movement toward the left or right side of the screen when ready (**Figure 1A**). Simultaneously, the monkey placed a wager on its choice by directing the saccade to either the upper target which represents a high bet, or the lower target which represents a low bet (**Figure 1A**). A high-bet correct choice would result in a larger juice reward than a low-bet correct choice. A high-bet incorrect choice would result in a time penalty (2-3 extra seconds of fixation required before the next stimulus presentation), whereas a low-bet incorrect choice was not penalized. Task difficulty was controlled by varying the percentage of coherently moving dots (motion strength), uniformly sampled from the following values: ± 0%, 3.2%, 6.4%, 12.8%, 25.6%, and 51.2%, where positive and negative represent rightward and leftward motion, respectively. Trials with 0% coherent motion were assigned a random direction and rewarded accordingly.

**Figure 1.**
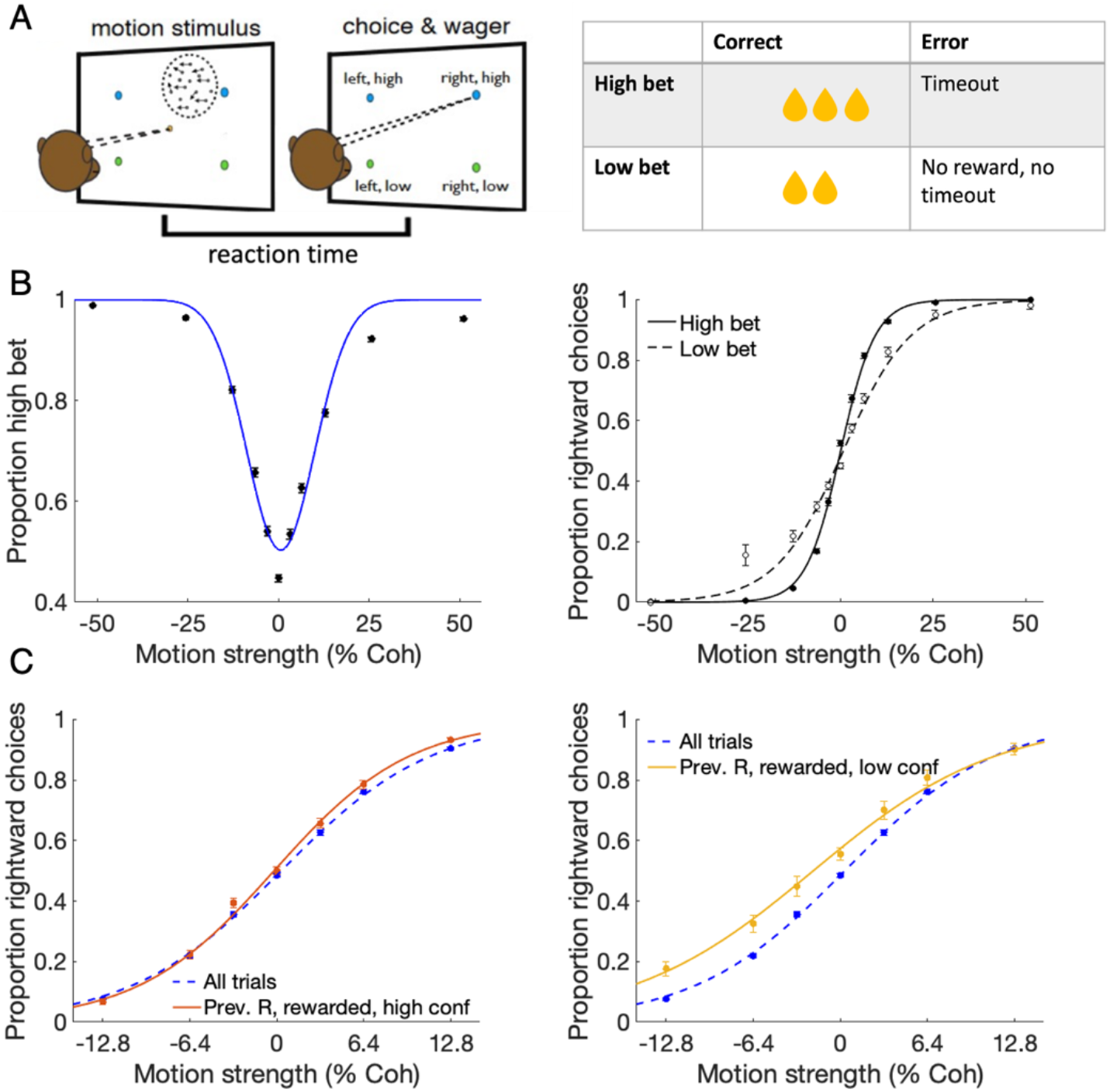
Task design and behavioral performance. (A). Random dot motion discrimination task with peri-decision wager. In each trial, the monkey views a dynamic random-dot stimulus and when ready reports its choice and wager with a single saccade to one of four targets. A high-bet choice will lead to a larger juice reward if correct, and a penalty timeout if incorrect. (B) Left: proportion of high-bet choices as a function of motion strength (% coherence; negative = leftward, positive = rightward. Smooth curve is a descriptive Gaussian fit. Right: psychometric functions plotted separately for high-bet and low-bet trials. (C) Psychometric functions over a narrower range of motion strengths to visualize the choice bias after rightward rewarded trials. Left panel shows the bias following a high-bet trial, right panel following a low-bet trial. Data are from monkey G (N = 35,157 trials) but both monkeys showed similar effects (see Fig. 2).

### Behavioral analysis

To quantify the monkeys’ bias as a function of trial history (**Fig. 2A,B**), we fit a simple logistic regression model to various subsets of trials defined by the previous trial’s motion strength, choice, wager, and outcome:

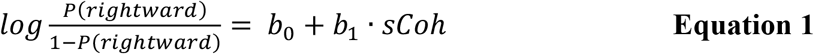

where *P(rightward)* is the proportion of trials ending in a rightward choice, *sCoh* is the signed motion strength on the current trial, and *b*_*i*_ are the regression coefficients. *b*_*0*_ measures the choice bias of the psychometric curve, with positive (negative) values indicating a rightward (leftward) choice bias. *b*_*1*_ measures the steepness, or sensitivity, of the psychometric curve.

**Figure 2.**
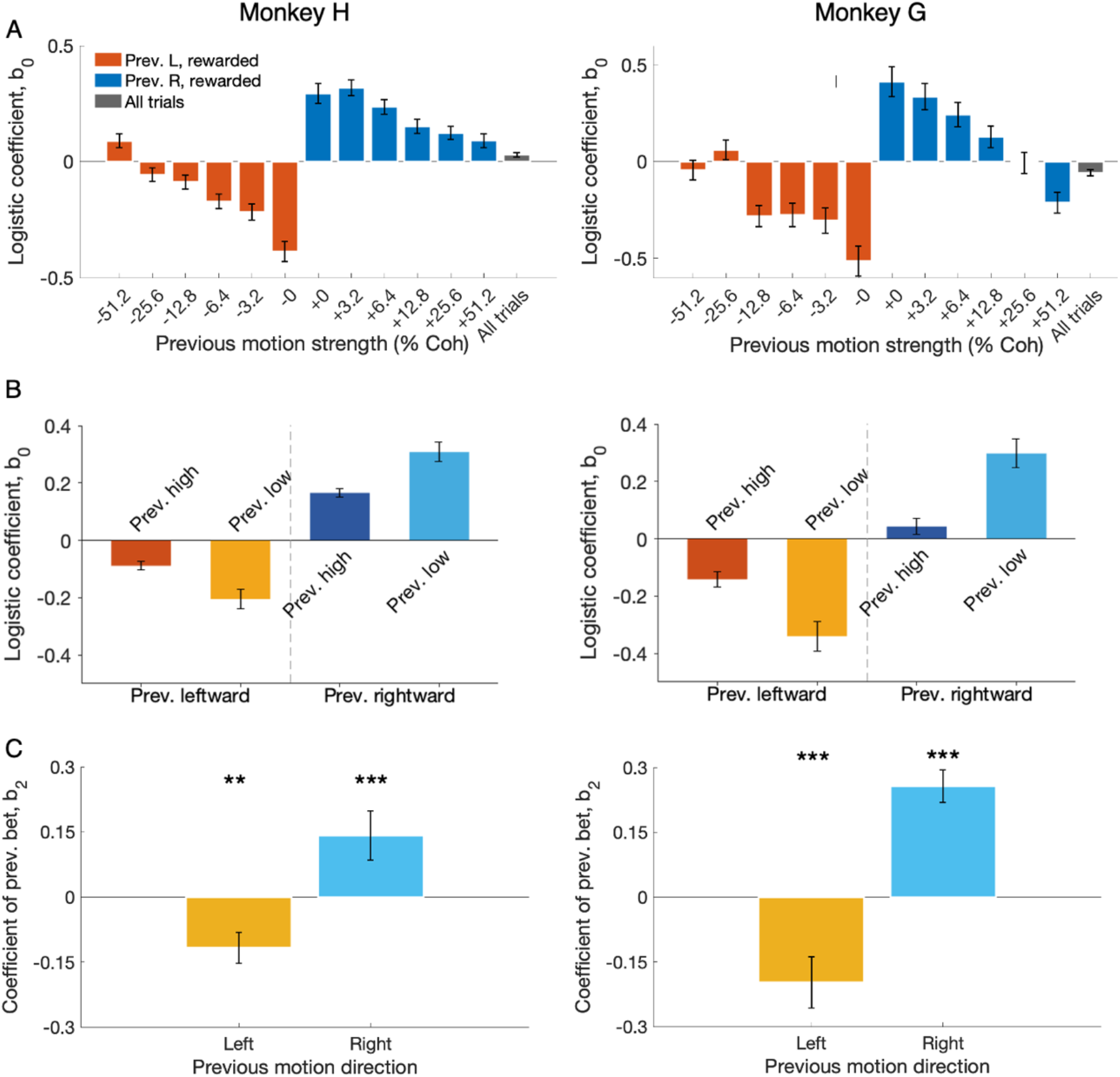
Win-stay bias is modulated by subjective confidence. (A) Logistic coefficient b_0_ (Eq. 1) as a function of the previous trial’s signed coherence. For zero-coherence trials, motion direction was assigned randomly. b_0_ quantifies the choice bias toward the previously rewarded side, with positive values indicating a rightward bias. Left: monkey H; right: monkey G. (B) Same data as in A but collapsed across motion strengths within each direction (orange/yellow: previous leftward trials; blue/cyan: previous rightward trials), and split by previous trial confidence (wager). (C) Logistic coefficient b_2_ (Eq. 2), separately for previous leftward and rightward choices. A statistically significant b_2_ indicates previous confidence has leverage over the current choice. The absolute value of b_2_ quantifies the bias of post-low-bet rewarded trials relative to post-high-bet rewarded trials. Left: monkey H; right: monkey G. **: p < 0.01; ***: p < 0.001.

To determine whether the confidence of the subject on the previous rewarded trial affects its current choice (**Fig. 2C**), a similar logistic regression model was fitted as above with the bet in the previous trial as an additional predictor:

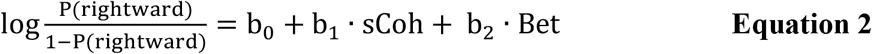

where *Bet* indicates the wager of the previous trial (0 = high bet, 1 = low bet) and *b*_*i*_ are the regression coefficients. *b*_*0*_ quantifies the choice bias, *b*_*1*_ tests for the effect of motion strength on choice, and *b*_*2*_ tests for the effect of previous-trial confidence on the choice bias. The magnitude of *b*_*2*_ quantifies the bias of a post-low-bet trial relative to a post-high-bet trial.

The effects of motion strength and previous outcome on choice accuracy were assessed with logistic regression (Purcell and Kiani, 2016):

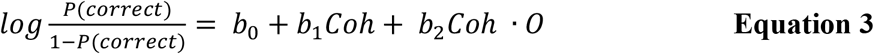

where *P(correct)* is the proportion of correct choices, *Coh* is the unsigned motion strength on the current trial, *O* is the outcome of the previous trial (0 = correct, 1 = incorrect) and *b*_*i*_ are the regression coefficients. *b*_*1*_ tests for the effect of motion strength on choice and *b*_*2*_ tests for the effect of the previous-trial outcome on the slope of the psychometric function. A significantly negative *b*_*2*_ indicates that the accuracy deteriorates following an error.

To analyze the relationship between previous confidence and subsequent choices while controlling for previous motion strength as a confounding factor, we performed an analysis of covariance (ANCOVA). The model included previous motion strength as a covariate, while previous confidence and current motion strength were treated as independent variables. The analysis was conducted separately for post-rightward-rewarded and post-leftward-rewarded trials. The ANCOVA model can be expressed as:

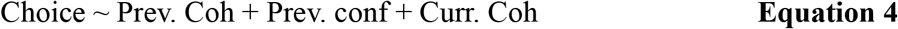

To determine how motion strength and the previous-trial outcome affect RT in the current trial, we used a linear regression model:

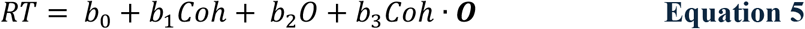

where *RT* is the RT on the current trial, *Coh* is the unsigned motion strength on the current trial, *O* is the outcome of the previous trial (0 = correct, 1 = incorrect), and *b*_*i*_ are the regression coefficients. *b*_*2*_ tests for the effect of the previous trial outcome on RT. *b*_*3*_ tests whether the extent of post-feedback adjustments of RT depend on the current motion strength.

To test for individual differences in the adjustments of RT to feedback, we constructed a mixed-effects linear regression model (**Equation 6**) with trials nested within the subjects:

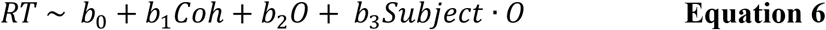

where *Subject* is the monkey identity.

To assess the effect of the previous trial confidence on RT, a linear regression model was used (**Equation 7**). For this analysis, we divided the trials according to whether the trial was post-correct or post-error and modeled them separately, to control for the effect of previous-trial feedback on RT. This was to ensure that the models would be easy to interpret.

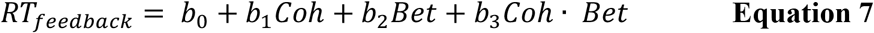

Here, *RT*_*feedback*_ is the RT on the current trial following an error or correct trial, *Bet* indicates the bet of the previous trial (0 = high and 1 = low for correct trials, whereas 0 = low and 1 = high for incorrect trials; this is to allow for easier interpretation of the coefficients) and *b*_*i*_ are the regression coefficients. *b*_*2*_ tests for the effect of previous-trial confidence on RT in the current trial. *b*_*3*_ determines whether the previous-trial confidence influences RT differently at different current motion strengths.

A mixed linear regression model was constructed to explain the individual differences between the monkeys’ strategies in adjusting RTs following an error or reward:

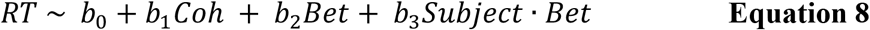

### Estimation of trial-by-trial choice bias

To quantify trial-by-trial choice bias, we implemented a logistic regression model based on the framework established by Hwang et al. (2017). In this model, the current choice is predicted by a weighted sum of multiple factors: the current stimulus, previous choice, previous outcome, and previous confidence rating. The weighted contribution of these trial history components provides an estimate of the internal choice bias that evolves from trial to trial.

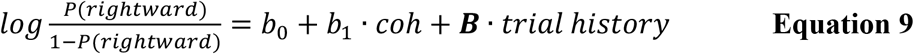

where *b*_*0*_ is a constant, *b*_*1*_ is weight for current coherence level, ***B*** is vector of weights for all the previous trial information.

### Modeling the motion task as a partially observable Markov decision process with temporal difference learning (POMDP-TD)

To model the effects of confidence on choice bias and post-feedback adjustments of RT, we constructed a belief-based RL model — the POMDP-TD model — leveraging the theory of sequential decision making under uncertainty combined with a temporal difference (TD) reinforcement learning algorithm (Khalvati et al., 2021; Lak et al., 2017; Lak, Okun, et al., 2020; Rao, 2010). In short, the agent continuously performs Bayesian updating of its belief over the true direction and strength of motion based on a stream of observations, termed momentary evidence, from the environment (**Figure 5**). The agent terminates the decision when the expected increase in confidence is less than the cost of continued observation. Upon termination, the choice is used to update the state-action values based on feedback using a TD algorithm. In this model, confidence-dependent choice bias arises from the across-trial updating of state-action values. The rate of evidence accumulation is scaled by the RPE of the previous trial to produce confidence-dependent post-feedback adjustments of RT.

Note that the POMDP algorithm was shown to have a one-to-one mapping to the parameters of the more commonly used drift-diffusion model (DDM), such that the latter can be seen as an implementation of the former (Khalvati et al., 2021). We chose to work within the POMDP framework for its flexibility and normativity (i.e., confidence as Bayesian belief), which seems a better match with the level of explanation embodied by the TD algorithm. Similar findings, absent the explicit confidence report, have been described within a DDM framework in previous work (Purcell & Kiani, 2016).

The hidden state of the environment for the POMDP-TD model is the signed coherence of the motion stimuli. The POMDP-TD model begins each trial with a prior belief about the signed coherence of the trial, b. = N(0, σ_0._). The prior distribution is approximated as a zero-mean Gaussian because the signed motion strengths used in this and other studies are concentrated near zero (Khalvati et al., 2021). At time step t of a trial with signed coherence c, the momentary observations, z_t_, will be drawn from a Gaussian distribution, N(c, w_z_) with mean equal to the signed coherence and variance equal to the true observation variance,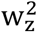. The true observation variance is unknown to the agent and hence the agent applies a learned observation variance, 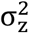, to construct its likelihood function, P(z |c) = N(z_t_, σ_z_).

With each observation, the agent performs a Bayesian update of its belief about the hidden state. From a Gaussian prior and a Gaussian likelihood function, the agent obtains a Gaussian posterior belief distribution (Khalvati et al., 2021) over the signed coherence:

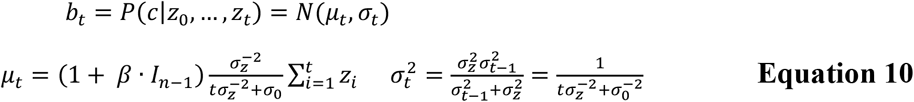

Note that, unlike previous work (Khalvati et al., 2021), the mean of the momentary observations (analogous to drift rate in the DDM) is modified by trial history (**Equation 10**), using an indicator variable (I_n−1_) for previous-trial outcome and confidence (**Equation 10**). β is a parameter, larger than 0 for tuning the impact of I_n−1_ on the current accumulation process. It is larger for low-confidence reward and high-confidence error. This modification allows trial history to impact RT in an outcome- and confidence-dependent manner, as observed in the data. Note that the model can reproduce post-error slowing (monkey H) or post-error speeding (monkey G) by allowing I_n−1_ to be negative or positive, respectively.

The agent’s provisional (‘online’) confidence at time step t, C_t_, is defined as the posterior probability of the direction with the greater posterior probability:

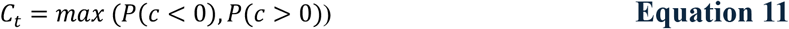

At each time step during the observing phase, the agent can either continue to observe or terminate the decision-making process and commit to a choice. A constant observation cost is assigned to collecting the momentary evidence (Rao, 2010). To terminate the observation optimally, the agent performs a one-step look-ahead search by comparing the expected increase in confidence to the cost of an observation (Khalvati et al., 2021). The agent halts new observations when the expected increase in its confidence is less than the cost of observation. To calculate the expected increase in confidence, the agent assumes that the next observation is sampled from the current belief distribution of the hidden state (Khalvati et al., 2021).

After terminating the observation phase, the agent uses a TD learning algorithm to make a choice (Lak et al., 2017; Rao, 2010). Since the reward only depends on choosing the correct direction, the model discretizes the continuous Gaussian belief distribution into two belief states, one state believing the motion direction is leftward, b_L_ and the other state believing the direction is rightward, b_R_:

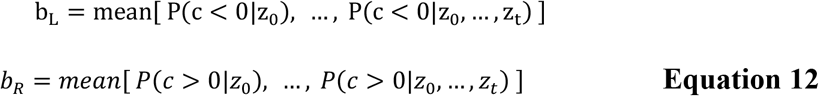

The model stores the state-action values of making a left (L) or right (R) choice given the belief state, resulting in four state-action pairs, Q (State = L or R, Action = L or R). On each trial, the action values, Q(L) and Q(R), are computed as the expected values of the state-action pairs:

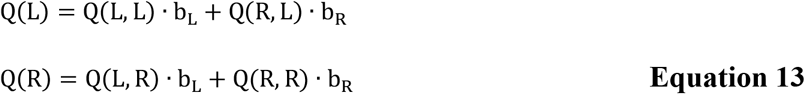

The choice is made by choosing the direction with the greater action value. The choice value, Q(choice), is given by:

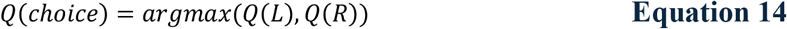

The decision confidence, C_c_, associated with the choice equals the belief state for that choice:

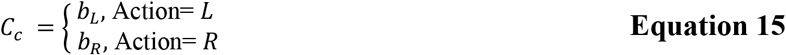

Upon receiving feedback (reward or punishment) from the environment, the model computes the RPE, δ, the difference between the reward and the choice value, integrating both past reward and the confidence about the accuracy of the current choice:

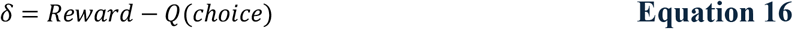

Lastly, the state-action pairs are updated using the RPE:

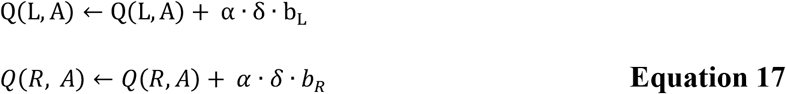

where α is the learning rate.

We simulated trials under this model with 150 agents performing the simplified version of the task described above. Each agent completed 2000 trials, allowing stable estimates of learning dynamics and choice behavior over time. Simulation results were obtained by averaging asymptotic behavior across agents, roughly after 300 trials with the learning rate used here. Parameters were hand-tuned to produce qualitative fits to key behavioral signatures in the monkey data, including asymptotic performance and the overall magnitude of post-feedback choice bias and RT adjustment. For the current study we present simulations rather than model fits because the goal was to demonstrate the capacity to reproduce the complex patterns of behavior and trial-history effects we observed. The quantitative similarity between simulated and empirical data demonstrates the feasibility of fitting the model, which can be used to compare with alternative models in future work.

### Neural analysis

#### Recording experiments

Detailed methods for neural data acquisition and preprocessing are described in the earlier report (Vivar-Lazo & Fetsch, 2026). Briefly, neural ensembles were recorded from the ventral subdivision of the lateral intraparietal area (LIPv, abbreviated as LIP) in the right hemisphere of two monkeys while they performed the peri-dw task. Recordings were obtained across 29 behavioral sessions (12 for monkey H, 17 for monkey G) using 32- or 128-channel probes (Diagnostic Biochips, Glen Burnie, MD). Single units (N=152) and multiunit clusters (N=225) (monkey H: single units 59, multi-units 148; monkey G: single units 93, multi-units 107) were sorted using Kilosort 2.0 combined with manual curation, and accepted for further analysis if their firing rate was significantly modulated during the delay period of a memory-saccade task.

#### Population decoding of history information

We conducted a population decoding analysis to examine neural signals related to previous choices and previous-trial confidence. For this we trained a support vector machine (SVM) with a radial basis function (RBF) kernel and L2 regularization, implemented using the Scikit-Learn library in Python.

The features consisted of firing rates from a simultaneously recorded neural ensemble in area LIP. The time-varying firing rate of each neural unit was quantified by counting the number of spikes in a sliding 200 ms window, with a step size of 50 ms. All units from the same recording session were utilized in decoding, provided they passed the acceptance criteria described above. Two SVM decoders were optimized to distinguish the neural activity on the current trial associated with the previous trial’s choice (‘left’ versus ‘right’) and previous trial’s confidence rating (‘high’ versus ‘low’), respectively.

The decoding process employed 10-fold cross-validation, where 90% of the data were used for training and 10% for testing. Decoding accuracy for each time point was calculated by averaging across all sessions (12 sessions for Monkey H and 17 sessions for Monkey G). The distance from the decoder’s decision boundary serves as a measure of the decoder’s prediction strength. We therefore utilized this distance at ± 0.95 s from stimulus motion onset to evaluate the degree of correlation between the neural encoding of trial history information and the estimated choice bias (**Fig. 8C-D**).

#### Estimating trial-by-trial decision variables

We implemented a separate SVM decoder to analyze neural dynamics related to the choice on the current trial. Similar to the history decoders detailed above, the current-trial decoder was trained using cross-validation to classify neural activity patterns associated with ‘leftward’ versus ‘rightward’ choices on the current trial. For each trial, we calculated a time-varying decision variable (DV), defined as the distance from the decoder’s decision boundary (Charlton & Goris, 2024; Kaufman et al., 2015; Kiani et al., 2014b). The time course of the DV (e.g., **Fig. 9**) represents the average population activity across trials, projected onto the dimension that best predicts the upcoming choice. To quantify temporal dynamics of the DV, we calculated its absolute value and averaged the magnitude across trials, segregating by current choice direction (left versus right) and conditioning on previous-trial choice and/or wager. We then performed a linear fit to the rising phase of the DV, specifically from its lowest point to its highest point after motion onset. Statistical differences in the decision variable’s slope, initial and final magnitudes across conditions were assessed using paired-sample t-tests. We conducted two separate analyses with different temporal alignments: one aligned to motion onset, and another aligned to saccade onset. This dual approach allowed us to examine decision-making dynamics from both stimulus presentation and response execution perspectives.

#### Experimental Design and Statistical Analysis

Two animals were used in the study as per field standards. Replication is implicit in the number of recorded neurons per animal per session, also consistent with field-specific conventions. Statistical tests, controls, corrections for multiple comparisons, and inclusion criteria are detailed for each separate analysis above.

## RESULTS

### Peri-decision wagering reflects confidence

In the peri-dw task (**Figure 1A**) monkeys report the net direction of motion (left or right) in a random-dot display, as well as a binary confidence judgment (high or low wager, or bet), by making a saccade to one of four targets. The monkey can indicate its choice and wager as soon as it is ready, so the task furnishes an estimate of decision time on every trial, and isolates the time window over which neural computations can support choice and confidence. A previous study using this task (Vivar-Lazo & Fetsch, 2026) analyzed behavior and neural activity on individual trials; here we focus on trial-history effects present in the same dataset. Both monkeys selected the high bet more frequently when motion strength was greater (**Figure 1B**, left). They also showed greater sensitivity (steeper psychometric functions) on high-bet vs. low-bet trials, meaning that the wager was correlated with accuracy even for a fixed stimulus strength (**Figure 1B**, right). This result held for identical repeats of the stimulus, modulo the RT (Vivar-Lazo & Fetsch, 2026). It implies that the monkeys did not wager solely based on an estimate of objective stimulus strength, but instead incorporated an internal prediction of accuracy or assessment of decision quality. Detailed relationships between wager frequency and response time (Vivar-Lazo & Fetsch, 2026) resembled human confidence judgments in similar tasks (Kiani et al., 2014a; van den Berg et al., 2016), further supporting the peri-dw assay as a valid measure of confidence in monkeys.

### Bias following reward (‘win-stay’) is modulated by confidence in the previous decision

We first evaluated whether positive feedback induces a choice bias on the following trial (**Figure 1C**) that scales with stimulus strength, as shown previously in other tasks and species (Lak et al., 2020). Figure 2A plots the bias derived from logistic regression (**Equation 1**, b_0_) as a function of the motion strength in the previous trial. A greater positive (negative) value of b_0_ indicates a greater rightward (leftward) bias on the current trial. Following a rewarded choice, the magnitude of b_0_ was greater when the motion strength in the previous trial was weaker, meaning that the animals displayed a greater tendency to repeat the rewarded choice as the previous trial became more difficult (**Figure 2A**). This shows that the win-stay bias in monkeys is dependent on stimulus strength, similar to earlier observations in rodents and humans (Lak, et al., 2020).

Because confidence is correlated with stimulus strength and (inversely) trial difficulty, previous studies attributed the stimulus strength-dependent bias to a learning rule that incorporates decision confidence, despite the absence of a behavioral confidence report. To test this interpretation more directly, we collapsed the data across motion strengths and plotted the post-reward bias (b_0_) split by whether the monkey bet high or low on the previous trial (**Figure 2B**), and indeed the magnitude of b_0_ was greater following a low bet (compare **Figure 1C** left vs. right to visualize the effect in terms of choice frequencies). To quantify the effect across directions, we constructed logistic regression models (**Equation 2**) separately for post-leftward-rewarded and post-rightward-rewarded trials, incorporating a coefficient for the previous bet (**Equation 2**, b_2_). We found that b_2_ was significantly different from zero (negative for previous left choice, positive for previous right choice) for both monkeys (**Figure 2C**), meaning they were more biased to repeat the previous rewarded choice when they reported lower confidence in the previous decision. Lastly, we split the data by the previous trial’s motion strength as well as wager, and found that the bias was generally greater following a low- vs. high-confidence rightward trial for each motion strength separately (**Figure 3A**, light blue bars tend to be higher than dark blue). The trend after a rewarded leftward choice was not as consistent (**Figure 3B**). To evaluate this pattern statistically, we constructed an ANCOVA model which quantifies the effect of previous-trial confidence on choices following a reward, controlling for the previous motion strength. This model showed a significant effect of previous confidence after a rightward-rewarded trial (**Supplementary Table 1.1**; F = 16.70, p ≪ 0.001), but not after a leftward-rewarded trial (**Supplementary Table 1.2**; F = 2.12, p = 0.15). In summary, monkeys showed a greater win-stay bias following a low-confidence rewarded choice, and this was not fully explained by the objective stimulus strength, suggesting that an internal sense of confidence gates the effect of feedback on the subsequent decision.

**Figure 3.**
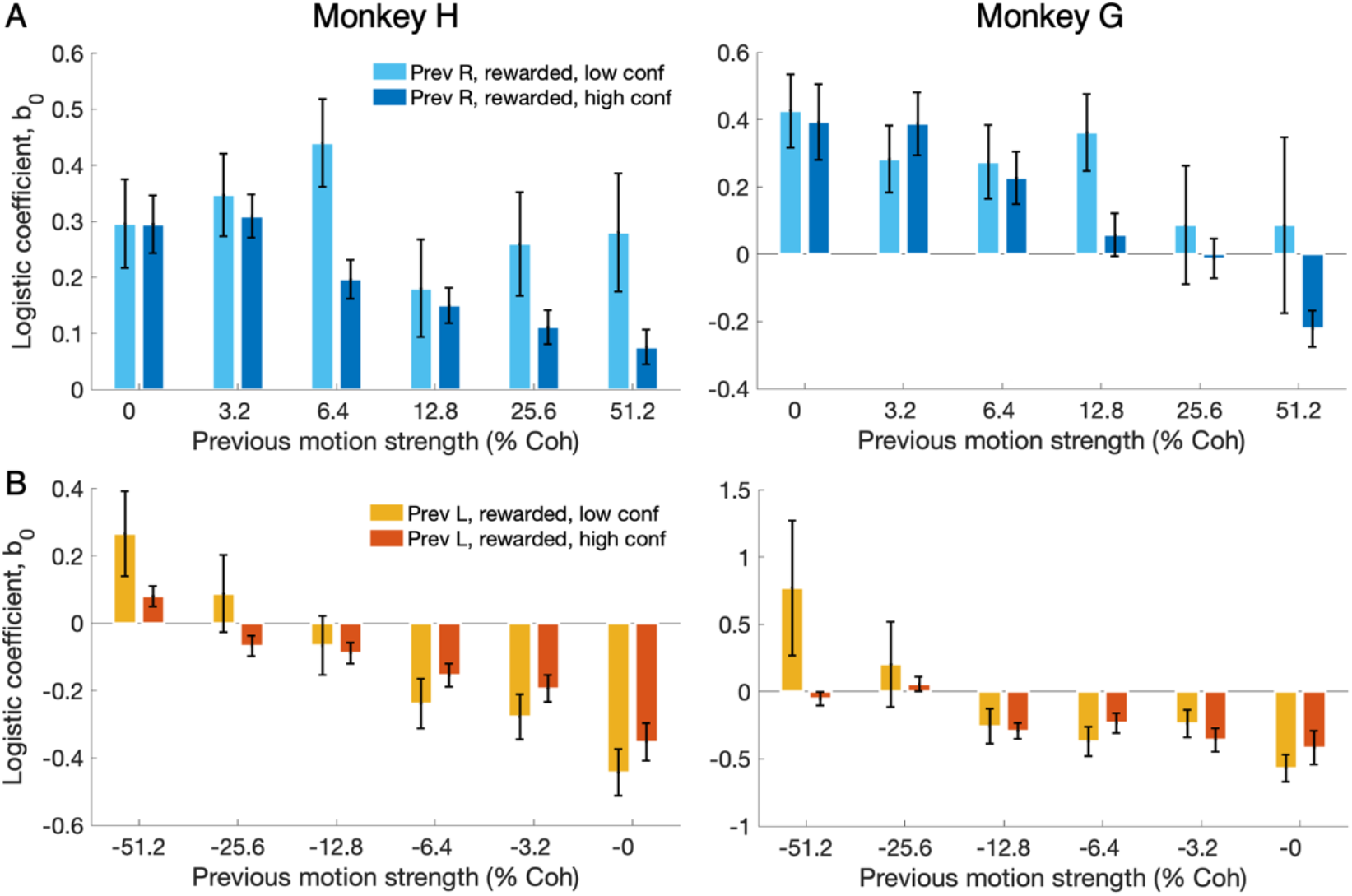
Choice bias is generally greater in post-reward, low-confidence trials within a given previous motion strength. (A) Choice bias on trials following a rightward, rewarded choice, separated by previous motion strength and wager (left: monkey H, right: monkey G). (B) Same as A but for previous leftward choices.

### Confidence predicts post-feedback adjustments of decision speed

In many kinds of decisions, response time provides a window on the policy of the decision maker and can therefore be a sensitive probe for trial-history effects. We found that monkeys showed significant changes in RT based on the outcome on the previous trial (**Equation 4**; Monkey H: b_2_ = 0.015 ± 0.002, p << 0.001; Monkey G: b_2_ = −0.0043 ± 0.006, p << 0.001), consistent with previous reports in humans and monkeys were found to slow down or speed up after negative feedback (Beatty et al., 2021; King et al., 2010; Purcell & Kiani, 2016). The direction and pattern of the effect differed by monkey (**Figure 4A; Equation 5**, b_3_ = 0.049 ± 0.004, t-test, p << 0.001): following an error, monkey H slowed down (**Figure 4A, left**), primarily at intermediate and high motion strengths, whereas monkey G sped up (**Figure 4A, right**) mostly on weak-motion trials. Post-error speeding is believed to occur when subjects emphasize speed over accuracy, perhaps ‘rushing through’ trials when aware of their reduced sensitivity (King et al., 2010; Purcell & Kiani, 2016). Consistent with this, monkey G showed a reduction in accuracy following an error (**Equation 3**, b_2_ = −1.45 ± 1.57, p = 0.01). Post-error slowing is assumed to play an adaptive role by allowing more evidence to accumulate before decision commitment, reducing the likelihood of an error (Botvinick et al., 2001; Holroyd et al., 2005; Laming, 1979). Contrary to this logic, monkey H was actually less accurate following an error, despite longer RTs (**Equation 3**, b_2_ = −3.52 ± 0.35, p << 0.001), as were several participants in previous studies. The lack of a simple relationship between RT and accuracy implies that post-feedback adjustments of RT might be caused by a mixture of changes in the decision-making process, including a reduction in sensitivity to incoming evidence that can decrease performance (Purcell & Kiani, 2016).

**Figure 4.**
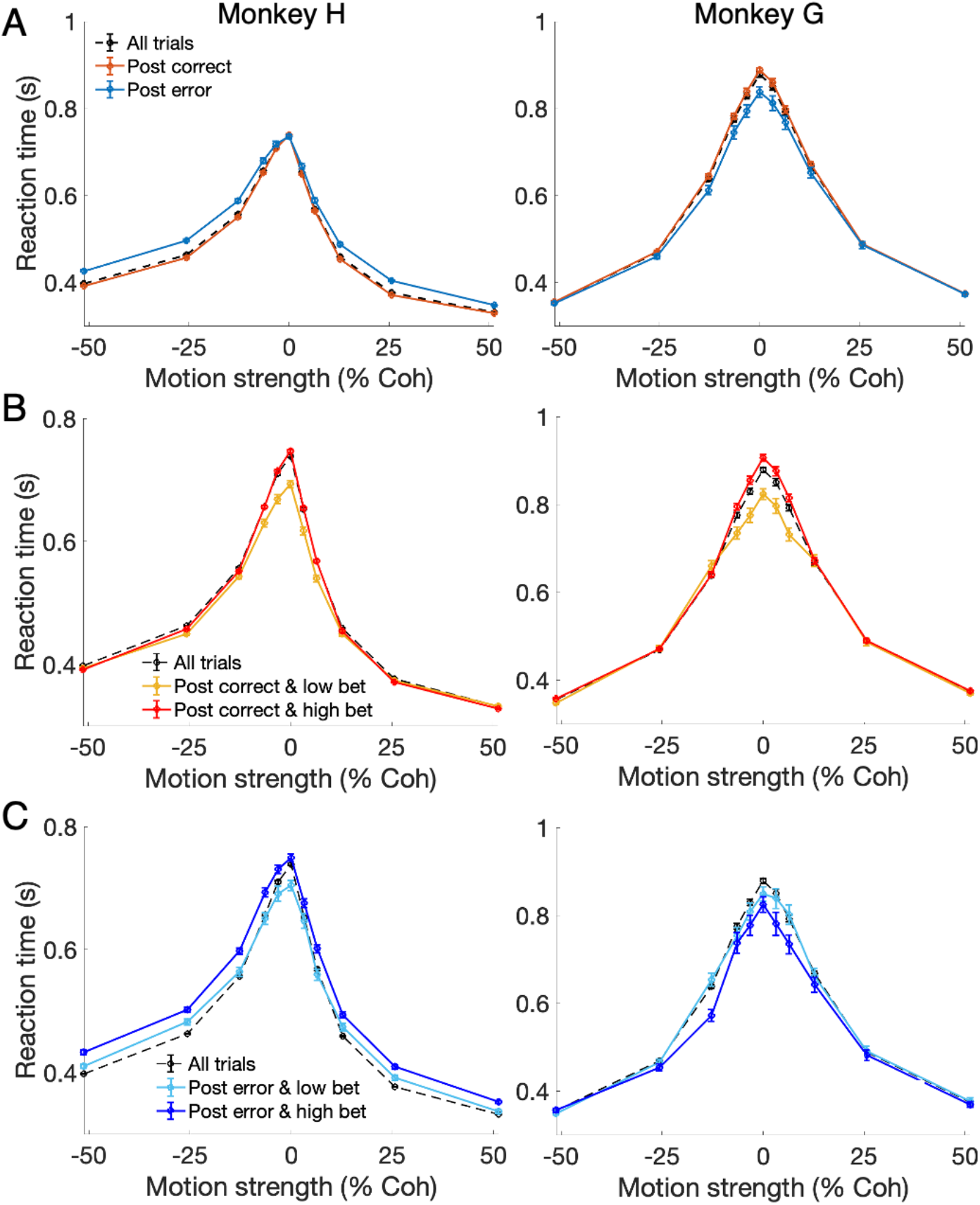
Effects of feedback and confidence on subsequent decision speed. (A) Reaction time (RT) as a function of motion strength, separated by previous trial outcome. Following an error, monkey H responded more slowly (left) whereas monkey G responded more quickly (right). (B) Post-correct RT separated by previous trial confidence. Both monkeys responded faster on the trial following a low-bet correct choice vs. a high-bet correct choice. (C) Post-error RT separated by previous trial confidence. Left: monkey H, who exhibited post-error slowing, slowed down more following a high-confidence error than a low-confidence error. Right: monkey G, who exhibited post-error speeding, reacted faster following a high-confidence error than a low-confidence error.

To determine whether confidence predicts post-feedback adjustment of RT, we constructed linear regression models separately for post-correct trials and post-error trials (**Equation 6**). After a rewarded trial, both monkeys showed faster RTs when the previous decision was rendered with low confidence (unexpected reward) compared to high confidence (expected reward; **Figure 4B; Equation 6**; Monkey H: b_2_ = −0.039 ± 0.004, p << 0.001; Monkey G: b_2_ = − 0.064 ± 0.006, p << 0.001). In contrast, the way in which confidence predicts post-error RT was different for the two monkeys (**Equation 7**, b_3_ = 0.058 ± 0.007, t-test, p << 0.001). Monkey H, who exhibited post-error slowing, had a significantly longer RT following a high-confidence error than a low-confidence error **(Figure 4C left; Equation 6**, b_2_ = 0.039 ± 0.004, p << 0.001). Monkey G, who exhibited post-error speeding, showed a significantly shorter RT following a high-confidence error than a low-confidence error **(Figure 4C right; Equation 6**, b_2_ = − 0.043 ± 0.010, p << 0.001). These results suggest that post-feedback adjustments of RT were dependent on subjects’ confidence, and challenge the view that such adjustments are shaped solely by a negative outcome. A less expected outcome—that is, a high-confidence error or low-confidence reward—induced a larger change in RT in the subsequent trial. These results provide evidence that the mismatch between actual and expected outcome drives post-feedback adjustment of RT (Holroyd et al., 2005) and that how much to update the subsequent decision policy (i.e. speed-accuracy trade-off) depends on confidence.

Interestingly, the changes in RT following a high-bet trial versus a low-bet trial were greatest when the current trial had a low motion strength (**Figure 4B, 4C; Equation 6**; Monkey H: post-correct b_3_ = 0.10 ± 0.01, p << 0.001, post-error b_3_ = −0.048 ± 0.018, p = 0.009; Monkey G: post-correct b_3_ = 0.15 ± 0.03, p << 0.001, post-error b_3_ = 0.093 ± 0.042, p = 0.028). This is consistent with the idea, common to many normative models, that when the current evidence is weak, prior expectations based on non-sensory factors (e.g., reward history, previous confidence, and outcome) will play a larger role in the current decision (Lak et al., 2017; Rao, 2010).

### POMDP-TD model reproduces key behavioral observations

To explain the observed behavioral patterns and build intuition about neural mechanisms, we constructed a hybrid model (**Figure 5**) that uses a partially observable Markov decision process (POMDP) to describe individual decisions (Khalvati et al., 2021) and a temporal difference reinforcement learning (TDRL) algorithm to capture changes in decision policy across trials. In this model, an RL agent processes noisy momentary evidence sampled from a Gaussian distribution determined by the current coherence level. On the time scale of individual trials, the agent employs Bayesian updating to continuously revise its belief distribution about the direction of motion, corresponding to the sign of the momentary evidence. This feature is consistent with the idea that a provisional or ‘online’ degree of confidence is maintained concurrently during decision formation (Balsdon et al., 2020; Dotan et al., 2018; Gherman & Philiastides, 2015; Vivar-Lazo & Fetsch, 2026). Across trials, the agent utilizes a TD learning algorithm specifically designed for partially observable environments which updates action values according to the agent’s confidence level.

**Figure 5.**
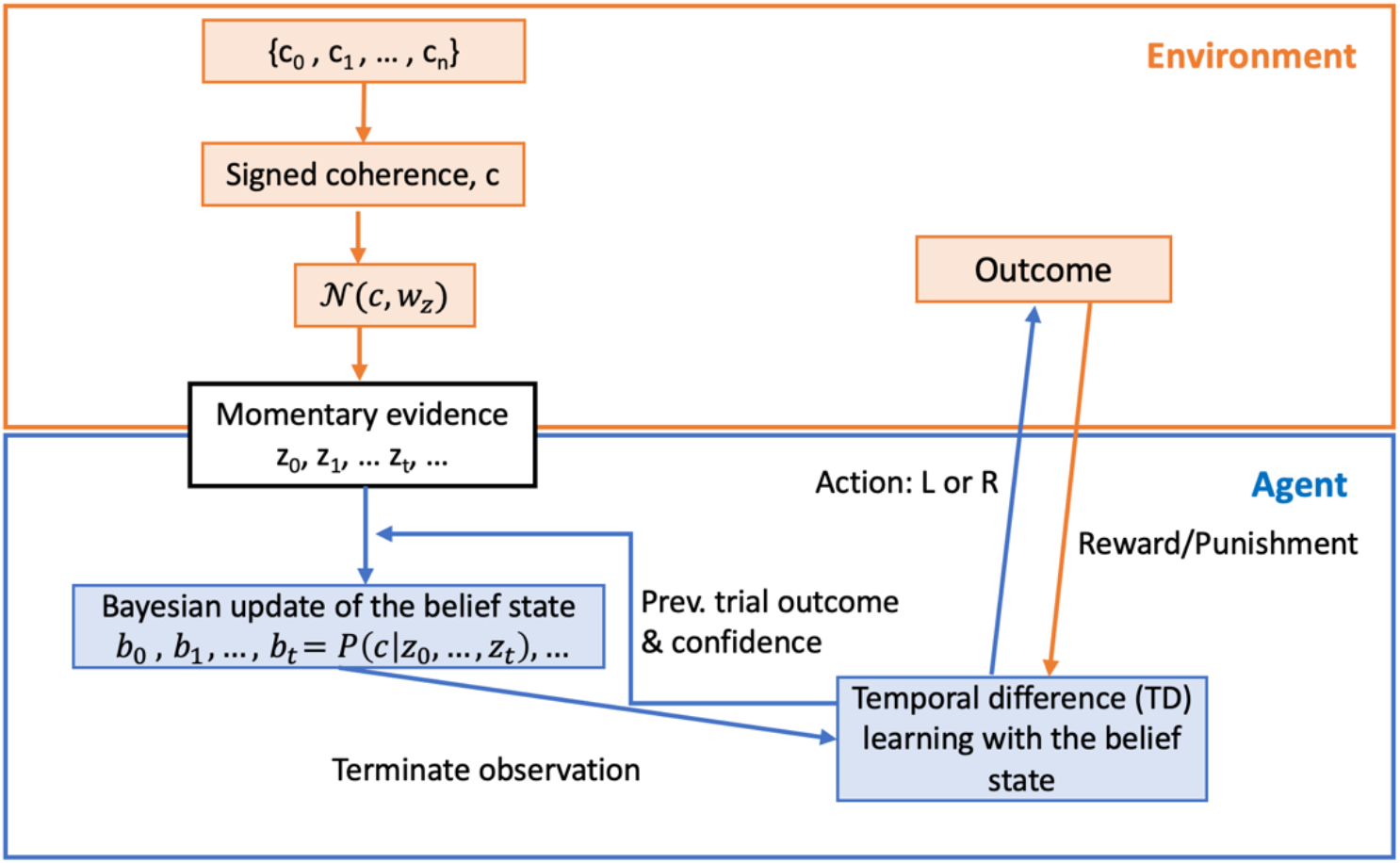
Schematic of the POMDP-TD model. The agent observes noisy momentary evidence and performs a Bayesian update of its belief distribution over all possible states (signed coherences, i.e. motion strengths and directions). This updating is influenced by previous-trial confidence and outcome. The agent terminates the observation using a one-step look-ahead algorithm by comparing the expected increase in confidence with the cost of observation, then makes a choice and updates its decision values based on feedback using a TD algorithm.

A key feature of the model is that both the confidence level and outcome from the previous trial serve as binary variables that influence subsequent evidence sampling. Model simulations reproduced the confidence-dependent choice bias and post-feedback adjustments of RT (**Figure 6**) that we observed behaviorally (**Figures 2-4**). As in the monkeys’ behavior, the model showed a decreasing choice bias with increasing motion strength (**Figure 6A**) and this bias was greater for previous low confidence trials (**Figure 6B**). Even when controlling for previous motion strength, the choice bias was greater following rewarded, low-confidence trials (**Figure 6C**). Beyond capturing the confidence-dependent choice bias, the model successfully reproduces RT adjustments observed in monkey behavior. The model can reproduce either post-error slowing (**Figure 6D, left**) or post-error speeding (**Figure 6E, left**) based on the sign of a single parameter (see **Eq. 10, Methods**). Similar to the monkeys, the magnitude of RT adjustment was greater when outcomes were less expected— specifically following high-confidence errors or low-confidence rewards (**Figure 6D,E, middle and right**).

**Figure 6.**
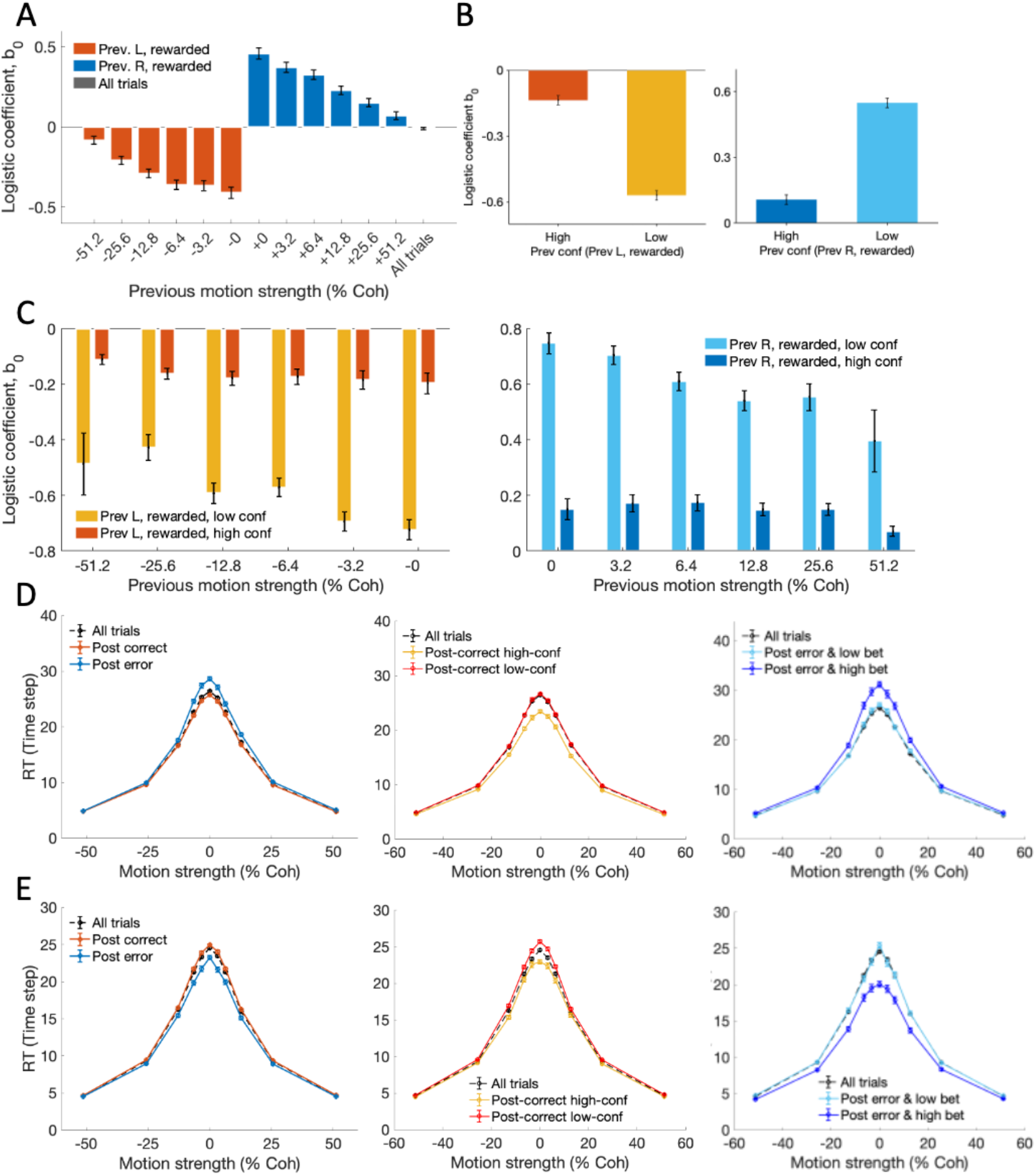
POMDP-TD model captures the confidence-dependent choice bias and post-feedback adjustment of RT. (A-C) Logistic regression analysis applied to simulated data from the model, in the same format as Figs. 2A, 2B, and 3, respectively. (D) Simulated RT functions separated by previous trial outcome & wager, showing confidence-dependent post-error slowing (compare to Fig. 4, left column (monkey H)). (E) Simulated RT functions for a separate instance of the model that exhibits confidence-dependent post-error speeding (compare to Fig. 4, right column (monkey G)).

In the POMDP-TD model, two key features account for the correspondence between simulated and observed behavior. First, the confidence-dependent choice bias is generated by the updating of state-action values through reward prediction error (RPE), which integrates both confidence and outcome information at the end of a trial. Second, the confidence-dependent adjustments in RT arise from a feedback loop where previous-trial confidence and outcome information directly influence the evidence accumulation process during the following observation phase. These mechanisms work together within the model’s POMDP framework to capture how perceptual decision-making dynamically incorporates trial history information. Moreover, the model’s architecture suggests that brain areas representing the current decision process should also encode trial history in a confidence-dependent manner. We therefore sought to test this hypothesis using recordings from area LIP.

### Neural correlates of confidence-dependent bias in LIP

Our previous study (Vivar-Lazo & Fetsch, 2026) found that LIP activity in the peri-dw task reflects a 2-dimensional DV that simultaneously resolves the choice and wager on the current trial. Conditioned on the previous trial’s wager, single-unit responses were heterogeneous (**Figure 7**). We found that 35% of neurons showed significant selectivity for previous-trial confidence (two-sample t-test, p<0.05) during the pre-stimulus period (−400 ms to 0 ms from stimulus onset) and 27% during the late post-stimulus period (200 ms to 400 ms from stimulus onset). These proportions were greater in monkey H (42% pre-stimulus and 32% late post-stimulus) than Monkey G (29% and 23%, respectively). Some neurons showed enhanced activity following low-confidence correct trials (**Figure 7A**, left), others exhibited suppressed activity under the same conditions (**Figure 7A**, middle), while a third group remained insensitive to the previous wager (**Figure 7A**, right). This heterogeneity in neuronal responses thus motivated us to examine how confidence-dependent history information might be encoded at the population level.

**Figure 7.**
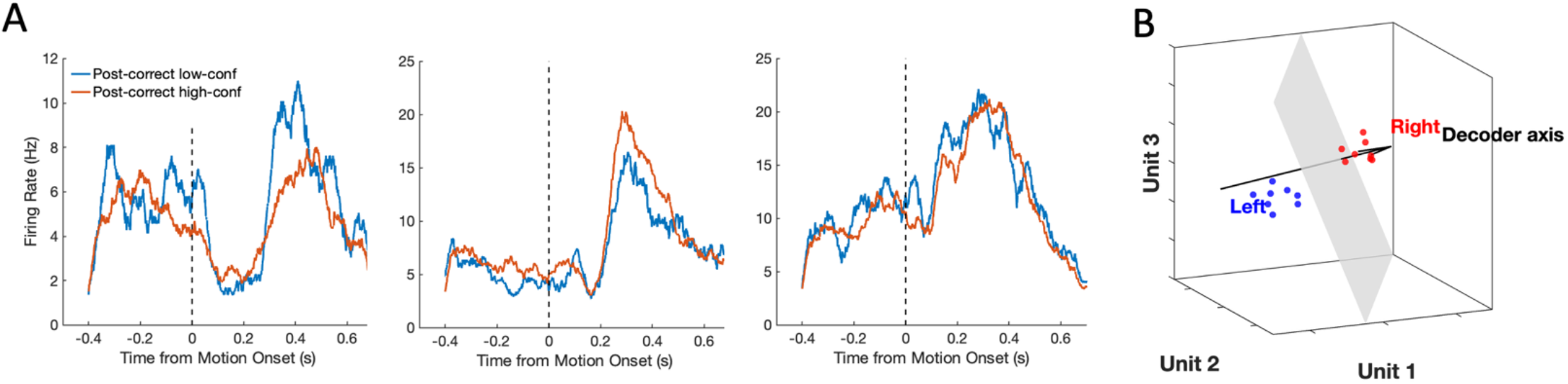
Heterogeneous trial-history signals in LIP. (A) Left: an example neuron that shows greater firing rate following low-confidence correct trials. Middle: example neuron that shows greater firing rate following high-confidence correct trials. Right: example neuron that does not distinguish previous trial confidence. (B) Schematic of population decoder (support vector machine) showing hypothetical neural activity in a simplified 3-dimensional space separated by a decision boundary.

We conducted a population decoding analysis (**Figure 7B**) to examine neural signals related to previous trial outcomes, using simultaneously recorded neurons from each individual session (mean = 14 units/session). Briefly, we trained support vector machines (SVMs) to classify neural activity patterns on a given trial that were associated with the previous trial’s choice (left vs. right) or wager (low vs. high). Because both monkeys exhibited a greater choice bias after a low-confidence correct trial, we compared cross-validated decoding accuracy for the previous outcome (correct trials only) conditioned on whether that choice was made with low vs. high confidence. Decoding accuracy for previous choice (**Figure 8A**) and previous wager (**Figure 8B**) was significantly above chance for an extended period surrounding the time of stimulus onset on the current trial, and was generally higher following a low-confidence rewarded trial compared to a high-confidence rewarded trial (Bonferroni-corrected p-values indicated by asterisks), except for previous choice in monkey G (**Figure 8A**, right). This implies that LIP encodes trial history information in a confidence-dependent manner, with an unexpected (low-confidence) reward leading to stronger encoding.

**Figure 8.**
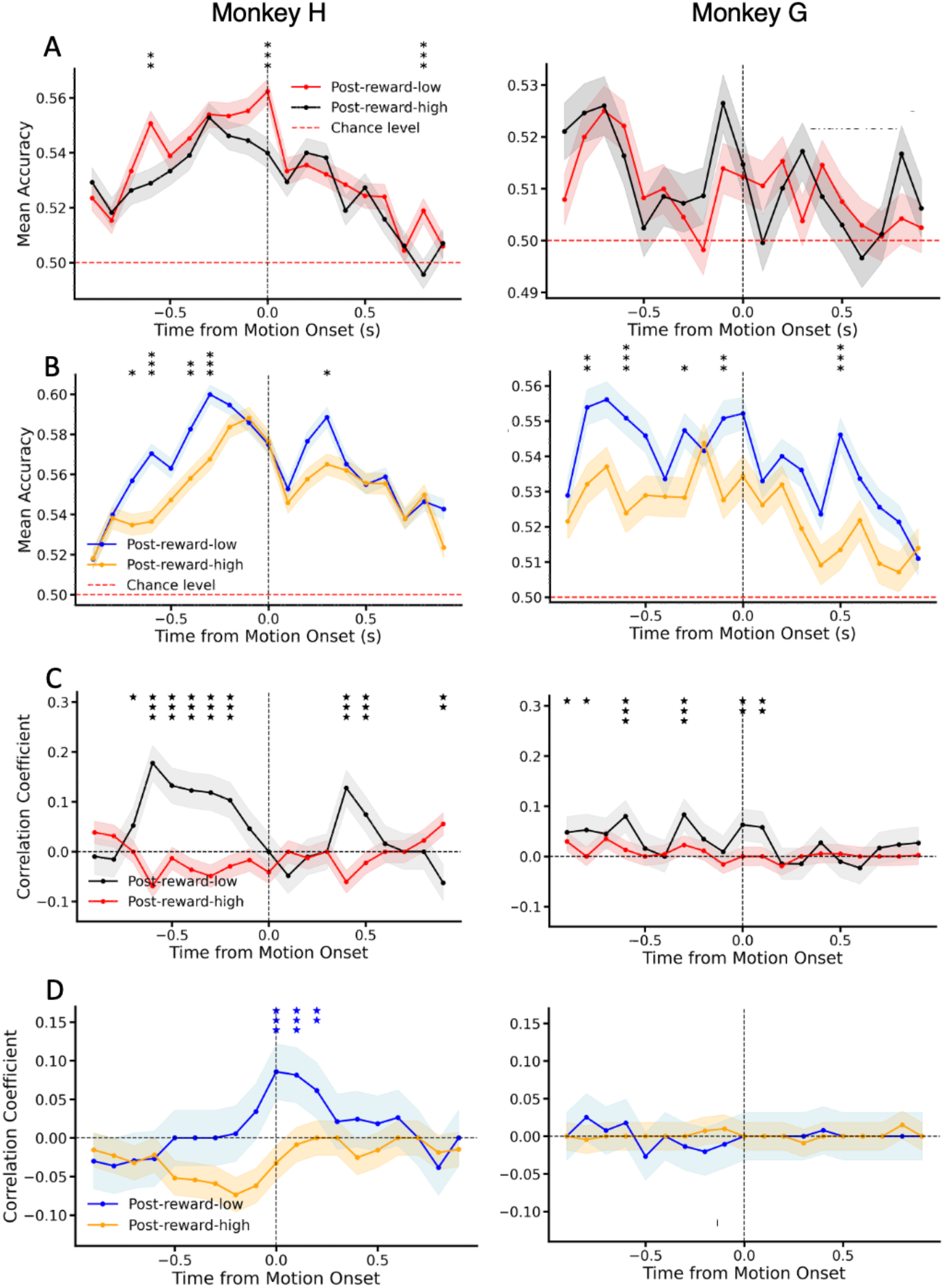
LIP populations encode trial history and indirectly reflect confidence-dependent bias. (A) Decoding accuracy of the previous choice (rewarded choices only) as a function of time from stimulus onset, separated by previous wager. (B) Same as A but for decoding of previous wager. (C) Correlation between a trial-by-trial estimate of the monkey’s bias and the strength of neural encoding of the previous choice, as a function of time relative to motion onset on the current trial. Asterisks indicate significant difference between the traces (*: p < 0.05; **: p < 0.01; ***: p < 0.001, Bonferroni corrected). (D) Same as C but for correlation between bias and encoding strength of previous confidence.

To test for a more direct link between trial-history signals and latent decision policy, we asked whether the encoding strength of trial history information was correlated with the monkey’s bias on a trial-by-trial basis. To estimate the bias, we constructed a linear regression model using the current stimulus and the immediate previous trial information, following the approach of Hwang et al. (2017) (**Equation 9**). Encoding strength of trial history was calculated as the distance between the N-dimensional neural population activity state and the decision boundary of the SVM classifier. We found a significant correlation between bias and encoding strength of the previous choice, but only after a low-confidence rewarded trial (**Figure 8C**; monkey H: p < 0.05 for 9 out of 19 time points; monkey G: p < 0.05 for 6 out of 19 time points, Bonferroni corrected). This effect was strongest before stimulus onset but persisted (or re-emerged) after stimulus onset, meaning it was present during decision formation on the current trial. For one monkey, there was also a positive correlation between bias and encoding strength for previous-trial confidence (**Figure 8D**, left). Together, these correlations support a functional coupling between the strength of trial-history representation in LIP and the magnitude of the animal’s choice bias. The next question is whether adjustments of decision speed might also be correlated with history-dependent changes in decision-related neural activity.

### Neural correlates of confidence-dependent adjustments of RT

To obtain a population-level representation of choice dynamics, we trained a separate SVM to predict the choice on the current trial, then used this decoder to extract a neural decision variable (DV) from the population activity (see Methods). Following previous approaches (Charlton & Goris, 2024; Kaufman et al., 2015; Kiani et al., 2014b; Peixoto et al., 2021) we defined the DV as the unsigned distance from the discriminant hyperplane of the choice decoder at each time point. Because LIP activity in the random-dot motion task has been shown to reflect a process of evidence accumulation (Churchland et al., 2011; Huk & Shadlen, 2005; Steinemann et al., 2024; Vivar-Lazo & Fetsch, 2026), the neural DV can be used to infer certain dynamic features of this process. We found that post-error adjustments of RT are reflected in the slope of the DV, assumed to reflect the rate of evidence accumulation. Monkey H, exhibiting post-error slowing showed a shallower slope for post-error trials (**Figure 9A, left**; t-test, p = 0.008; **Table 1**), whereas Monkey G, displaying post-error speeding (**Figure 9A, right**; t-test, p=0.038; **Table 1**). In contrast, the starting and ending points of the DV conditioned on different trial outcomes were not significantly different (**Figure 9A** and **Supplementary Figure 1A; Table 4**), suggesting that post-error slowing/speeding is mediated by rate of accumulation rather than a change in the initial state or termination bound of the decision process (cf. Purcell & Kiani, 2016).

**Figure 9.**
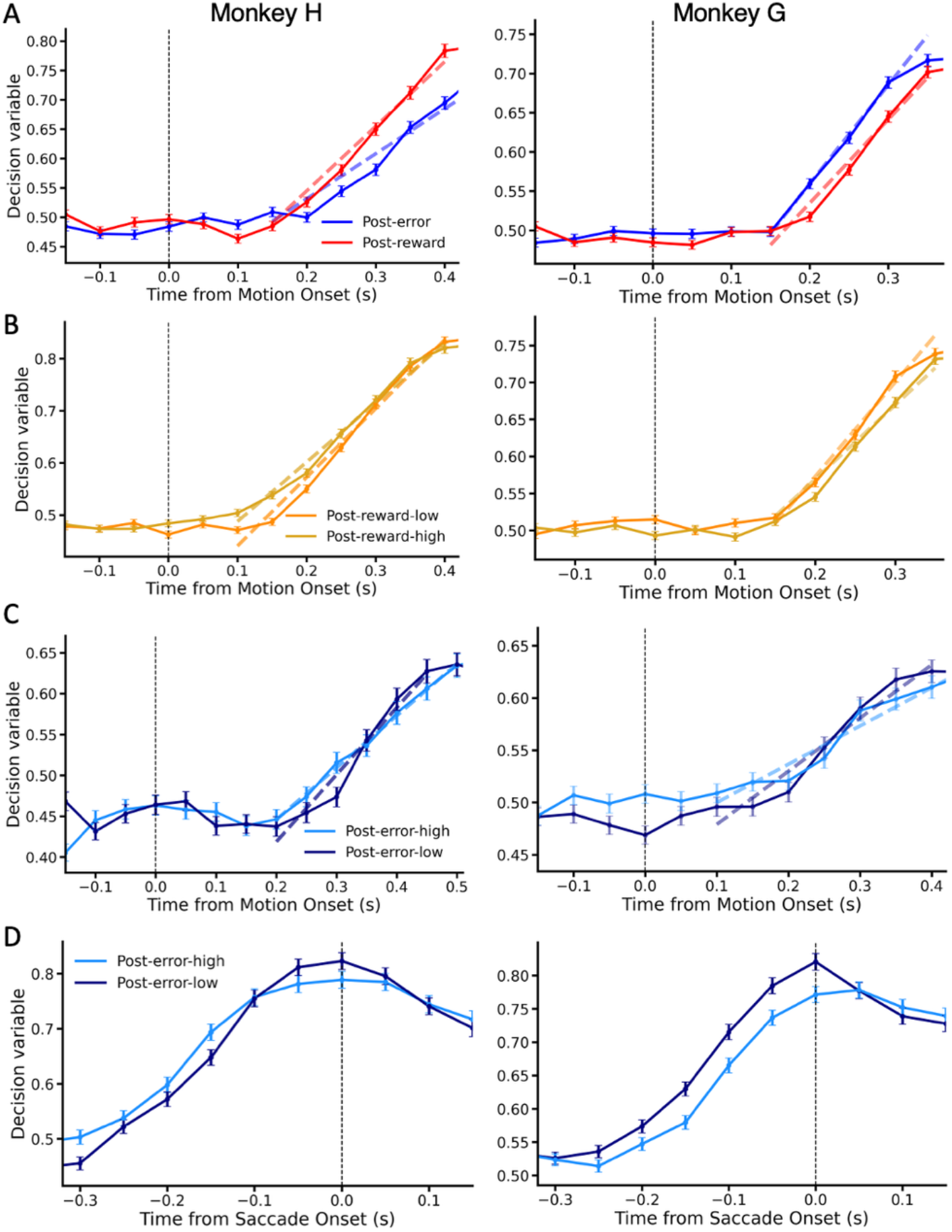
Dynamics of neural population activity in LIP reflects post-feedback adjustments of RT. Left column = monkey H, right column = monkey G. (A) Average time course of the neural DV (prediction strength of choice decoder) separated by whether the previous trial was correct (rewarded) or incorrect. Dashed lines indicate a linear fit to the data points during the ramping phase of the DV (from its lowest point to its highest point after motion onset). (B) Same as A but for post-reward trials only, split by previous trial confidence (wager). (C) Average DV traces for trials following an error, separated by previous trial confidence (wager), aligned to motion onset. (D) Same as C but aligned to saccade onset.

**Table 1.** DV slopes following correct vs. error trials.

|  | Trial type | Slope | Constant | R-squared | Slope p-value |
| --- | --- | --- | --- | --- | --- |
| Monkey H | Post-error | $0.7669 \pm 0.0820$ | 0.3783 | 0.9356 | 0.0075 |
| | Post-reward | $1.0430 \pm 0.0790$ | 0.3405 | 0.9721 | |
| Monkey G | Post-error | $1.2598 \pm 0.0369$ | 0.3075 | 0.9983 | 0.0380 |
| | Post-reward | $1.0656 \pm 0.1031$ | 0.3217 | 0.9727 | |

Further analysis revealed that these slope effects were modulated by previous-trial confidence. The DV slope was steeper for both monkeys following a low-confidence correct trial (**Figure 9B; Table 2**: t-test, Monkey H: p = 0.036; Monkey G: p = 0.018), correlating with faster RT after low-confidence correct trials (i.e., an unexpected reward) (**Figure 4B)**. Recall that the two monkeys showed distinct patterns in their post-error behavior that were further modulated by confidence: Monkey H showed enhanced post-error slowing following high-confidence errors, while Monkey G showed enhanced post-error speeding after high-confidence errors (**Figure 4C**). Despite these opposing behavioral patterns, decoded LIP activity in both monkeys revealed shallower DV slopes following high-confidence error trials (**Figure 9C; Table 3**: t-test, Monkey H: p = 0.015; Monkey G: p = 0.013), suggesting that unexpected errors led to decreased rates of evidence accumulation. This dissociation also implies that differences in DV slope do not automatically or trivially follow from differences in RT.

**Table 2.** DV slopes following high vs. low confidence correct trials.

|  | Condition | Slope | Constant | R-squared | Slope p-value |
| --- | --- | --- | --- | --- | --- |
| Monkey H | Post-reward low | $1.3188 \pm 0.0799$ | 0.3084 | 0.9820 | 0.0356 |
| | Post-reward high | $1.1397 \pm 0.0589$ | 0.3733 | 0.9868 | |
| Monkey G | Post-reward low | $1.2739 \pm 0.0978$ | 0.3181 | 0.9883 | 0.0184 |
| | Post-reward high | $1.0014 \pm 0.0864$ | 0.3691 | 0.9711 | |

**Table 3.**
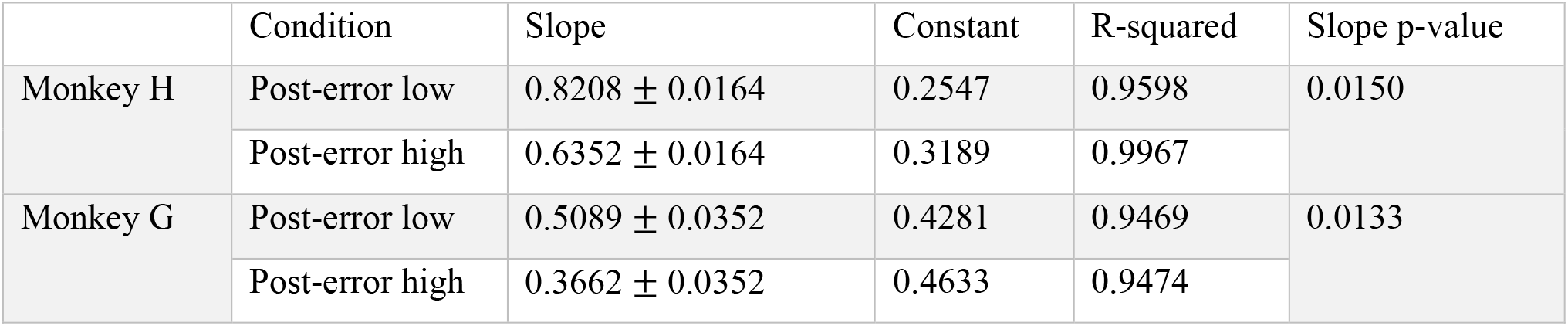
DV slopes following high vs. low confidence error trials.

|  | Condition | Slope | Constant | R-squared | Slope p-value |
| --- | --- | --- | --- | --- | --- |
| Monkey H | Post-error low | $0.8208 \pm 0.0164$ | 0.2547 | 0.9598 | 0.0150 |
| | Post-error high | $0.6352 \pm 0.0164$ | 0.3189 | 0.9967 | |
| Monkey G | Post-error low | $0.5089 \pm 0.0352$ | 0.4281 | 0.9469 | 0.0133 |
| | Post-error high | $0.3662 \pm 0.0352$ | 0.4633 | 0.9474 | |

In Monkey G, who exhibited post-error speeding, the DV showed two distinct characteristics following high-confidence errors: an elevated starting point and a reduced end point (**Figure 9C & D** right; **Table 4:** t-test, Monkey G: starting point: p < 0.001; end point: p = 0.002). This pattern reveals a complex interaction where two mechanisms — start point elevation and early termination of evidence accumulation — appear to offset the expected slowing from decreased evidence accumulation rate. In contrast, Monkey H showed no significant differences in the starting and ending points of the DV (**Figure 9C & D** left; **Table 4**: t-test, starting point: p = 0.957; end point: p = 0.057). Instead, this monkey’s enhanced post-error slowing after high-confidence error trials is straightforwardly explained by a shallower slope of the DV in these trials. Taken together, the findings demonstrate that LIP population activity integrates confidence and reward information from the previous trial to adjust the evidence accumulation process (especially its rate) in the current trial. These neural correlates are strikingly predictive of idiosyncratic behavioral patterns in the two animals, and are broadly consistent with a computational framework for within- and across-trial decision processes as embodied by the POMDP-TD model.

**Table 4.** Comparison of DV starting and end points across trial outcomes.

|  | Condition | Starting pt | p-value | End pt | p-value |
| --- | --- | --- | --- | --- | --- |
| Monkey H | Post-error low | $0.463 \pm 0.317$ | 0.9567 | $0.823 \pm 0.411$ | p = 0.0574 |
| | Post-error high | $0.464 \pm 0.322$ | | $0.789 \pm 0.413$ | |
| Monkey G | Post-error low | $0.469 \pm 0.321$ | p < 0.001 | $0.821 \pm 0.468$ | p = 0.0022 |
| | Post-error high | $0.508 \pm 0.339$ | | $0.771 \pm 0.442$ | |

## DISCUSSION

Adaptive behavior in uncertain environments requires not only learning what actions to pursue given a particular world state (i.e., a value function), but also adjusting how to go about ascertaining the current world state (i.e., the perceptual decision process). Here we investigated how confidence—the subjective belief in the correctness of a decision—shapes this aspect of learning, capitalizing on a recently developed paradigm for nonhuman primates that measures choice, RT, and confidence on every behavioral trial (Vivar-Lazo & Fetsch, 2026). We found that trial-history effects on choice bias and RT were modulated by the monkey’s self-reported confidence on the previous trial, and that confidence-dependent policy adjustments have modest but significant neural correlates in parietal area LIP. The behavioral effects were recapitulated by a reinforcement learning model in which the belief state at the time of a decision carries forward to influence both the accumulation process and action values on the next trial, suggesting predictions that can be tested in future experiments.

Our work was enabled by the fact that perceptual decisions are invariably influenced by previous outcomes, even in tasks where trials are designed to be independent. These influences can be complex (Roy et al., 2021; Zhu & Kuchibhotla, 2024), though here we focus on two of the simplest examples: the win-stay bias, where subjects repeat previously rewarded choices (Lee et al., 2004), and post-error slowing or speeding (Beatty et al., 2021; Laming, 1979; Purcell & Kiani, 2016). Such behaviors are objectively suboptimal because the reward is based only on the stimulus evidence in the current trial. From the point of view of the organism, however, they could reflect a normative strategy that evolved for environments in which states tend to change more slowly than in a randomized laboratory experiment. Indeed the ubiquity of these effects, and the fact that extensive training fails to eliminate them (Gold et al., 2008; Mochol et al., 2021), suggests they may be deeply embedded in the neural mechanisms underlying choice behavior.

A key observation is that history effects depend systematically on task difficulty or stimulus strength. Studies in mice, rats, and humans have shown that win-stay biases are greatest immediately after a trial with weak sensory evidence (Lak et al., 2020; Mendonça et al., 2020). Similarly, post-error adjustments of RT are larger following more difficult trials (Purcell & Kiani, 2016). Because confidence is correlated with stimulus strength, these results have been used to claim that confidence gates the influence of feedback on subsequent choices (Braun et al., 2018; Lak, Hueske, et al., 2020). However, confidence reflects more than objective stimulus strength (Odegaard et al., 2018), and can be decoupled from accuracy in a variety of ways (Del Cul et al., 2009; Drugowitsch et al., 2014; Rahnev et al., 2012), so its empirical link to trial-history effects has thus far been indirect. Some previous studies used proxies of confidence based on the relationship between stimulus strength and accuracy under assumptions of signal-detection theory (SDT; Braun et al., 2018) or a more general formulation known as statistical decision confidence (SDC; Hangya et al., 2016; Lak et al., 2017). These frameworks are useful and well-supported, but they fail to robustly explain metacognitive sensitivity (Boundy-Singer et al., 2023; Shekhar & Rahnev, 2021), and cannot account for temporal factors (Desender et al., 2021; Kiani et al., 2014a). Notably, several recent studies demonstrate that the canonical prediction of SDT/SDC—the ‘folded-X’ pattern of confidence as a function of stimulus strength, conditioned on accuracy—is violated under many conditions (Desender et al., 2021; Hellmann et al., 2023; Kiani et al., 2014a; Vivar-Lazo & Fetsch, 2026; Xue et al., 2026), implying that it cannot be uncritically used to identify a proxy or correlate of confidence. By measuring confidence behaviorally on each trial, we address this issue and show that trial-history effects are modulated by subjective confidence even when controlling for stimulus difficulty. This validates the prevailing hypothesis and also helps constrain the underlying neural mechanism. Specifically, it implicates cognitive signals that incorporate internal noise, including metacognitive noise (Boundy-Singer et al., 2023; Shekhar & Rahnev, 2021), not merely analysis of stimulus features that correlate with difficulty and hence confidence.

Although higher-order representations of confidence play a dominant role in metacognition research (for reviews see Fleming, 2024; Goris et al., 2025), a growing body of work indicates that confidence can be decoded from the same sensorimotor areas engaged in forming the primary decision (Balsdon et al., 2020; Balsdon & Philiastides, 2024; Dotan et al., 2018; Dou et al., 2024; Gherman & Philiastides, 2015; Vivar-Lazo & Fetsch, 2026). This parallel computation of a ‘sensorimotor’ form of confidence raises intriguing possibilities for local implementation of adaptive learning mechanisms.

Using population analyses of spiking activity in area LIP—a region associated with evidence accumulation (Steinemann et al., 2024) and decision termination (Stine et al., 2023)—we found that the strength of encoding of previous choices was correlated with a trial-by-trial estimate of choice bias (**Figure 8**). This correlation suggests a direct link between neural representations of past decisions and behavioral policy adjustments, indicating that the same populations computing current choices may also store and utilize information from previous trials in a confidence-dependent manner. Similar observations were made by Hwang and colleagues (2017) who found that mouse PPC activity during the inter-trial interval of a visual decision task predicted the animal’s trial-by-trial choice bias. When PPC was inactivated, the behavioral choice-history effects diminished, suggesting that PPC is critical for mediating the effect of trial history on the current decision (see also Akrami et al., 2018). In monkeys, a substantial fraction of LIP neurons was found to represent action value signals derived from previous outcomes in a reinforcement learning task (Seo et al., 2009).

Taken together, these results and the current findings implicate PPC, along with prefrontal cortex and basal ganglia, in the network implementing outcome-based modifications to decision policy. The dynamics of our decoded decision variable suggest a potential mechanism for one such modification. We observed systematic changes in the ramping slope of the DV (presumed to reflect the rate of evidence accumulation) following unexpected outcomes, with adjusted slopes after both low-confidence rewards and high-confidence errors (**Figure 9**). Similarly, Purcell & Kiani (2016) showed that LIP activity in monkeys was influenced by errors in a manner that could explain post-error adjustments of RT. Following a negative outcome, LIP dynamics reflected a weaker urgency signal and decreased rate of evidence accumulation, consistent with the behavioral observation of post-error slowing (PES). These predictive relationships between neural signatures and individual behavioral patterns suggest that confidence-dependent learning involves specific modifications of the evidence accumulation process, implemented by a network that includes LIP.

Because confidence-dependent learning entails a comparison between predicted and actual outcomes, its underlying mechanism seems likely to overlap with circuits computing reward prediction error (RPE). Classic work demonstrates that midbrain dopamine neurons encode RPE (Schultz et al., 1997), including in a temporally extended fashion during a random-dot motion task (Nomoto et al., 2010). Lak and colleagues reanalyzed these data to show that macaque dopamine neuron activity is consistent with a representation of confidence (Lak et al., 2017), and subsequently reported that both the ventral tegmental area and medial prefrontal cortex of mice encode confidence-related variables and influence subsequent choices (Lak, Okun, et al., 2020). However, it remains unclear whether decision confidence per se, rather than a correlate like stimulus strength or perceived difficulty, shapes dopaminergic RPE signals. It would be exciting for future studies to uncover a pathway by which cortical representations of confidence are relayed to mesolimbic or nigrostriatal circuitry in order to influence the RPE calculation. Then to close the loop, one might test whether dopaminergic projections to PPC itself (Lewis et al., 2001; van Kempen et al., 2022)—in addition to their better-known targets in prefrontal cortex—provide the mechanistic link between metacognitive assessment and adaptive alterations of the decision process.

## Conclusion

Our findings confirm that subjective confidence serves as an important gate for learning from experience, enabling adaptive adjustments of decision policy based on the reliability of internal decision signals. The encoding of confidence-dependent trial history within decision circuits including LIP raises the possibility of local mechanisms that could enable rapid behavioral adaptation. A hybrid POMDP-TD model reproduces the behavioral patterns across our two animal subjects, and offers a framework for generating normative hypotheses about the biological implementation of reinforcement learning under perceptual uncertainty.

## Acknowledgments

The authors thank Ofelia Garalde for animal care, Bill Nash, Bill Quinlan, and Dean Carpenter for technical assistance. This work was supported by the National Institute of Neurological Disorders and Stroke (RF1NS132910) and the Whitehall Foundation (#2021-05-112). C.R.F. is also supported by the France-Merrick Foundation. Current address for MVL: Dept. of Neuroscience & Zuckerman Institute, Columbia University, New York, NY, USA, 10027.

**Supplementary Figure 1.**
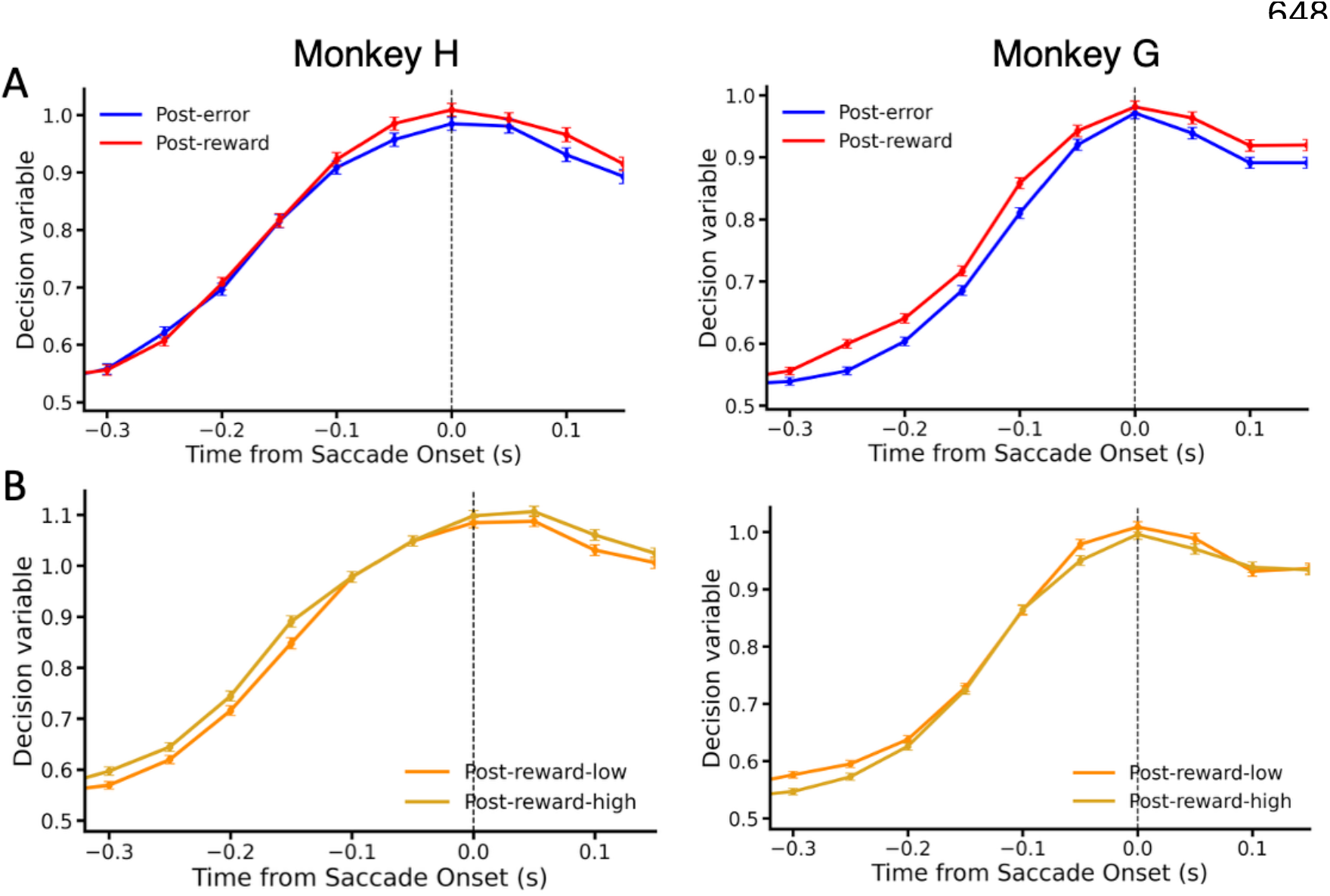
End points of the neural DV, shown aligned to saccade onset, do not differ as a function of previous trial outcome or confidence. (A) Conditioned on previous correct vs error trials; (B) Conditioned on previous confidence following correct trials. Left: monkey H (post-error slowing monkey). Right: monkey G (post-error speeding monkey).

**Supplementary Table 1.** ANCOVA: Effect of previous confidence on post-rewarded choices, controlling for the previous motion strength. **Table 1.1**. post-rightward-rewarded trials across the two monkeys. **Table 1.2**. post-leftward-rewarded trials across the two monkeys.

| **Supplementary Table 1.1** | **Post-rewarded-post-rightward choice ~ Prev. %Coh + Prev. conf + Curr. %Coh** |  |  |  |
| --- | --- | --- | --- | --- |
|  | Sum Sq | df | F value | p-value |
| Prev. % Coh (Covariate) | 5.7 | 1 | 38.78 | 4.77e-10 *** |
| Prev. conf. | 2.4 | 1 | 16.70 | 4.38e-05 *** |
| Curr. % Coh | 6978.0 | 1 | 47749.35 | < 2.2e-16 *** |
| **Supplementary Table 1.2** | **Post-rewarded-post-leftward choice ~ Prev. %Coh + Prev. bet + Curr. %Coh** |  |  |  |
|  | Sum Sq | df | F value | p-value |
| Prev. % Coh (Covariate) | 11.4 | 1 | 78.08 | < 2.0e-16 *** |
| Prev. conf. | 0.3 | 1 | 2.12 | 0.15 |
| Curr. % Coh | 7053.6 | 1 | 48313.03 | < 2.0e-16 *** |

## References

Akrami, A., Kopec, C. D., Diamond, M. E., & Brody, C. D. (2018). Posterior parietal cortex represents sensory history and mediates its effects on behaviour. Nature, 554(7692), 368–372. 10.1038/nature25510

Balsdon, T., & Philiastides, M. G. (2024). Confidence control for efficient behaviour in dynamic environments. Nature Communications, 15(1), 9089. 10.1038/s41467-024-53312-3

Balsdon, T., Wyart, V., & Mamassian, P. (2020). Confidence controls perceptual evidence accumulation. Nature Communications, 11(1). 10.1038/s41467-020-15561-w

Beatty, P. J., Buzzell, G. A., Roberts, D. M., Voloshyna, Y., & McDonald, C. G. (2021). Subthreshold error corrections predict adaptive post-error compensations. Psychophysiology, 58(6), e13803. 10.1111/PSYP.13803

Botvinick, M. M., Braver, T. S., Barch, D. M., Carter, C. S., & Cohen, J. D. (2001). Conflict Monitoring and Cognitive Control. Psychological Review, 108(3), 624–652. 10.1037/0033-295X.I08.3.624

Boundy-Singer, Z. M., Ziemba, C. M., & Goris, R. L. T. (2023). Confidence reflects a noisy decision reliability estimate. Nature Human Behaviour, 7(1), 142–154. 10.1038/s41562-022-01464-x

Braun, A., Urai, A. E., & Donner, T. H. (2018). Adaptive History Biases Result from Confidence-Weighted Accumulation of past Choices. Journal of Neuroscience, 38(10), 2418–2429. 10.1523/JNEUROSCI.2189-17.2017

Charlton, J. A., & Goris, R. L. T. (2024). Abstract deliberation by visuomotor neurons in prefrontal cortex. Nature Neuroscience, 27(6), 1167–1175. 10.1038/s41593-024-01635-1

Churchland, A. K., Kiani, R., Chaudhuri, R., Wang, X. J., Pouget, A., & Shadlen, M. N. (2011). Variance as a Signature of Neural Computations during Decision Making. Neuron, 69(4), 818–831. 10.1016/j.neuron.2010.12.037

Crapse, T. B., Lau, H. & Basso, M. A. (2018) A Role for the Superior Colliculus in Decision Criteria. Neuron 97, 181-194.e6

Del Cul, A., Dehaene, S., Reyes, P., Bravo, E., & Slachevsky, A. (2009). Causal role of prefrontal cortex in the threshold for access to consciousness. Brain, 132(9), 2531–2540. 10.1093/brain/awp111

Desender, K., Boldt, A., Verguts, T., & Donner, T. H. (2019). Confidence predicts speed-accuracy tradeoff for subsequent decisions. ELife, 8. 10.7554/eLife.43499

Desender, K., Ridderinkhof, K. R., & Murphy, P. R. (2021). Understanding neural signals of post-decisional performance monitoring: An integrative review. ELife, 10. 10.7554/ELIFE.67556

Dotan, D., Meyniel, F., & Dehaene, S. (2018). On-line confidence monitoring during decision making. Cognition, 171, 112–121. 10.1016/j.cognition.2017.11.001

Dou, W., Martinez Arango, L. J., Castaneda, O. G., Arellano, L., Mcintyre, E., Yballa, C., & Samaha, J. (2024). Neural Signatures of Evidence Accumulation Encode Subjective Perceptual Confidence Independent of Performance. Psychological Science, 35(7), 760–779. 10.1177/09567976241246561

Drugowitsch, J., Mendonça, A. G., Mainen, Z. F., & Pouget, A. (2019). Learning optimal decisions with confidence. Proceedings of the National Academy of Sciences, 116(49), 24872–24880. 10.1073/pnas.1906787116

Drugowitsch, J., Moreno-Bote, R., & Pouget, A. (2014). Relation between Belief and Performance in Perceptual Decision Making. PLoS ONE, 9(5), e96511. 10.1371/journal.pone.0096511

Fleming, S. M. (2024). Metacognition and Confidence: A Review and Synthesis. Annual Review of Psychology, 75(1), 241–268. 10.1146/annurev-psych-022423-032425

Frömer, R., Nassar, M. R., Bruckner, R., Stürmer, B., Sommer, W., & Yeung, N. (2021) Response-based outcome predictions and confidence regulate feedback processing and learning. eLife 10, e62825 (2021).

Gherman, S., & Philiastides, M. G. (2015). Neural representations of confidence emerge from the process of decision formation during perceptual choices. NeuroImage, 106, 134–143. 10.1016/j.neuroimage.2014.11.036

Gold, J. I., Law, C. T., Connolly, P., & Bennur, S. (2008). The relative influences of priors and sensory evidence on an oculomotor decision variable during perceptual learning. Journal of Neurophysiology, 100(5), 2653–2668. 10.1152/JN.90629.2008

Gold, J. I., & Shadlen, M. N. (2007). The neural basis of decision making. In Annual Review of Neuroscience (Vol. 30, pp. 535–574). 10.1146/annurev.neuro.29.051605.113038

Goris, R. L. T., Fu, Z., & Fetsch, C. R. (2025). Computational and Neuronal Basis of Visual Confidence. Annual Review of Vision Science, 11(1), 385–410. 10.1146/annurev-vision-110323-120909

Hangya, B., Sanders, J. I., & Kepecs, A. (2016). A Mathematical Framework for Statistical Decision Confidence. Neural Computation, 28(9), 1840–1858. 10.1162/NECO_a_00864

Hellmann, S., Zehetleitner, M., & Rausch, M. (2023). Simultaneous modeling of choice, confidence, and response time in visual perception. Psychological Review, 130(6), 1521–1543. 10.1037/rev0000411

Holroyd, C. B., Yeung, N., Coles, M. G. H., & Cohen, J. D. (2005). A mechanism for error detection in speeded response time tasks. Journal of Experimental Psychology: General, 134(2), 163–191. 10.1037/0096-3445.134.2.163

Huk, A. C., & Shadlen, M. N. (2005). Neural activity in macaque parietal cortex reflects temporal integration of visual motion signals during perceptual decision making. Journal of Neuroscience, 25(45), 10420–10436. 10.1523/JNEUROSCI.4684-04.2005

Hwang, E. J., Dahlen, J. E., Mukundan, M., & Komiyama, T. (2017). History-based action selection bias in posterior parietal cortex. Nature Communications, 8(1), 1–14. 10.1038/s41467-017-01356-z

Kaufman, M. T., Churchland, M. M., Ryu, S. I., & Shenoy, K. V. (2015). Vacillation, indecision and hesitation in moment-by-moment decoding of monkey motor cortex. ELife, 4. 10.7554/eLife.04677

Khalvati, K., Kiani, R., & Rao, R. P. N. (2021). Bayesian inference with incomplete knowledge explains perceptual confidence and its deviations from accuracy. Nature Communications, 12(1), 5704. 10.1038/s41467-021-25419-4

Kiani, R., Corthell, L., & Shadlen, M. N. (2014a). Choice Certainty Is Informed by Both Evidence and Decision Time. Neuron, 84(6), 1329–1342. 10.1016/j.neuron.2014.12.015

Kiani, R., Cueva, C. J., Reppas, J. B., & Newsome, W. T. (2014b). Dynamics of Neural Population Responses in Prefrontal Cortex Indicate Changes of Mind on Single Trials. Current Biology, 24(13), 1542–1547. 10.1016/J.CUB.2014.05.049

Kiani, R., & Shadlen, M. N. (2009). Representation of Confidence Associated with a Decision by Neurons in the Parietal Cortex. Science, 324(5928), 759–764. https://www.jstor.org/stable/20493893

King, J. A., Korb, F. M., Von Cramon, D. Y., & Ullsperger, M. (2010). Post-Error Behavioral Adjustments Are Facilitated by Activation and Suppression of Task-Relevant and Task-Irrelevant Information Processing. Journal of Neuroscience, 30(38), 12759–12769. 10.1523/JNEUROSCI.3274-10.2010

Lak, A., Hueske, E., Hirokawa, J., Masset, P., Ott, T., Urai, A. E., Donner, T. H., Carandini, M., Tonegawa, S., Uchida, N., & Kepecs, A. (2020). Reinforcement biases subsequent perceptual decisions when confidence is low, a widespread behavioral phenomenon. ELife, 9. 10.7554/eLife.49834

Lak, A., Nomoto, K., Keramati, M., Sakagami, M., & Kepecs, A. (2017). Midbrain Dopamine Neurons Signal Belief in Choice Accuracy during a Perceptual Decision. Current Biology, 27(6), 821–832. 10.1016/J.CUB.2017.02.026/ATTACHMENT/47F01226-F596-4939-8F51-F2271C0F25E3/MMC1.PDF

Lak, A., Okun, M., Moss, M. M., Gurnani, H., Farrell, K., Wells, M. J., Reddy, C. B., Kepecs, A., Harris, K. D., & Carandini, M. (2020). Dopaminergic and Prefrontal Basis of Learning from Sensory Confidence and Reward Value. Neuron, 105(4), 700-711.e6. 10.1016/J.NEURON.2019.11.018/ATTACHMENT/364E7724-D88B-42D1-89F5-7C5203E5ADB4/MMC1.PDF

Laming, D. (1979). Choice reaction performance following an error. Acta Psychologica, 43(3), 199–224. 10.1016/0001-6918(79)90026-X

Lee, D., Conroy, M. L., McGreevy, B. P., & Barraclough, D. J. (2004). Reinforcement learning and decision making in monkeys during a competitive game. Cognitive Brain Research, 22(1), 45–58. 10.1016/j.cogbrainres.2004.07.007

Lewis, D. A., Melchitzky, D. S., Sesack, S. R., Whitehead, R. E., Auh, S., & Sampson, A. (2001). Dopamine transporter immunoreactivity in monkey cerebral cortex: Regional, laminar, and ultrastructural localization. Journal of Comparative Neurology, 432(1), 119–136. 10.1002/cne.1092

Mendonça, A. G., Drugowitsch, J., Vicente, M. I., DeWitt, E. E. J., Pouget, A., & Mainen, Z. F. (2020). The impact of learning on perceptual decisions and its implication for speed-accuracy tradeoffs. Nature Communications, 11(1), 1–15. 10.1038/s41467-020-16196-7

Mochol, G., Kiani, R., & Moreno-Bote, R. (2021). Prefrontal cortex represents heuristics that shape choice bias and its integration into future behavior. Current Biology, 31(6), 1234-1244.e6. 10.1016/J.CUB.2021.01.068

Morcos, A. S., & Harvey, C. D. (2016). History-dependent variability in population dynamics during evidence accumulation in cortex. Nature Neuroscience, 19(12), 1672–1681. 10.1038/nn.4403

Newsome, W., & Pare, E. (1988). A selective impairment of motion perception following lesions of the middle temporal visual area (MT). Journal of Neuroscience, 8(6), 2201–2211. 10.1523/JNEUROSCI.08-06-02201.1988

Nomoto, K., Schultz, W., Watanabe, T. & Sakagami, M. (2010). Temporally Extended Dopamine Responses to Perceptually Demanding Reward-Predictive Stimuli. Journal of Neuroscience, 30, 10692–10702.

Odegaard, B., Grimaldi, P., Cho, S. H., Peters, M. A. K., Lau, H., & Basso, M. A. (2018). Superior colliculus neuronal ensemble activity signals optimal rather than subjective confidence. Proceedings of the National Academy of Sciences of the United States of America, 115(7), E1588–E1597. 10.1073/pnas.1711628115

Peixoto, D., Verhein, J. R., Kiani, R., Kao, J. C., Nuyujukian, P., Chandrasekaran, C., Brown, J., Fong, S., Ryu, S. I., Shenoy, K. V., & Newsome, W. T. (2021). Decoding and perturbing decision states in real time. Nature, 591(7851), 604–609. 10.1038/s41586-020-03181-9

Purcell, B. A., & Kiani, R. (2016). Neural Mechanisms of Post-error Adjustments of Decision Policy in Parietal Cortex. Neuron, 89(3), 658–671. 10.1016/j.neuron.2015.12.027

Rahnev, D. A., Maniscalco, B., Luber, B., Lau, H., & Lisanby, S. H. (2012). Direct injection of noise to the visual cortex decreases accuracy but increases decision confidence. Journal of Neurophysiology, 107(6), 1556–1563. 10.1152/jn.00985.2011

Rao, R. P. N. (2010). Decision Making Under Uncertainty: A Neural Model Based on Partially Observable Markov Decision Processes. Frontiers in Computational Neuroscience, 4, 146. 10.3389/fncom.2010.00146

Roy, N. A., Bak, J. H., Akrami, A., Brody, C. D., & Pillow, J. W. (2021). Extracting the dynamics of behavior in sensory decision-making experiments. Neuron, 109(4), 597-610.e6. 10.1016/j.neuron.2020.12.004

Schultz, W., Dayan, P., & Montague, P. R. (1997). A Neural Substrate of Prediction and Reward. Science, 275(5306), 1593–1599. 10.1126/science.275.5306.1593

Seo, H., Barraclough, D. J., & Lee, D. (2009). Lateral Intraparietal Cortex and Reinforcement Learning during a Mixed-Strategy Game. Journal of Neuroscience, 29(22), 7278–7289. 10.1523/JNEUROSCI.1479-09.2009

Shadlen, M. N., & Newsome, W. T. (2001). Neural Basis of a Perceptual Decision in the Parietal Cortex (Area LIP) of the Rhesus Monkey. Journal of Neurophysiology, 86(4), 1916–1936. 10.1152/jn.2001.86.4.1916

Shekhar, M., & Rahnev, D. (2021). The nature of metacognitive inefficiency in perceptual decision making. Psychological Review, 128(1), 45–70. 10.1037/rev0000249

Steinemann, N. A., Stine, G. M., Trautmann, E. M., Zylberberg, A., Wolpert, D. M., & Shadlen, M. N. (2024). Direct observation of the neural computations underlying a single decision. ELife. 10.7554/eLife.90859.2

Stine, G. M., Trautmann, E. M., Jeurissen, D., & Shadlen, M. N. (2023). A neural mechanism for terminating decisions. Neuron, 111(16), 2601-2613.e5. 10.1016/j.neuron.2023.05.028

Urai, A. E., Braun, A., & Donner, T. H. (2017). Pupil-linked arousal is driven by decision uncertainty and alters serial choice bias. Nature Communications, 8(1), 1–11. 10.1038/ncomms14637

van den Berg, R., Zylberberg, A., Kiani, R., Shadlen, M. N., & Wolpert, D. M. (2016). Confidence Is the Bridge between Multi-stage Decisions. Current Biology, 26(23), 3157–3168. 10.1016/j.cub.2016.10.021

van Kempen, J., Brandt, C., Distler, C., Bellgrove, M. A., & Thiele, A. (2022). Dopamine influences attentional rate modulation in Macaque posterior parietal cortex. Scientific Reports, 12(1). 10.1038/s41598-022-10634-w

Vivar-Lazo, M., & Fetsch, C. R. (2026). Neural basis of concurrent deliberation toward a choice and confidence judgment. Nature Neuroscience, 29(1), 159–170. 10.1038/s41593-025-02116-9

Xue, K., Fung, H., & Rahnev, D. (2026). Stimulus reliability but not boundary distance manipulations violate the folded-X pattern of confidence. Cognition, 272, 106490. 10.1016/j.cognition.2026.106490

Zhu, Z., & Kuchibhotla, K. V. (2024). Performance errors during rodent learning reflect a dynamic choice strategy. Current Biology, 34(10), 2107-2117.e5. 10.1016/j.cub.2024.04.017

